# Proton pump inhibitor-associated microbiomes shape systemic host responses

**DOI:** 10.64898/2026.11.10.765563

**Authors:** Karlis A. Moors, Saskia Weber-Stiehl, Hannah Jebens, Bram Nap, Daniel Fässler, Ameneh Mehrjerd, Martina A. Guggeis, Silke S Heinzmann, Alesia Walker, Corinna Bang, Fenja Amrei Schuran, Georgios Marinos, Kristina Schlicht, Jennifer Hörlöck, Jan Taubenheim, Andre Franke, Wolfgang Lieb, Matthias Laudes, Almut Heinken, Philippe Schmitt-Kopplin, Philip Rosenstiel, Johannes Hertel, Ines Thiele, Mathieu Groussin, Mathilde Poyet, Felix Sommer, Christoph Kaleta

**Affiliations:** Institute of Experimental Medicine, Kiel University, Kiel, Germany; Institute of Clinical Molecular Biology, Kiel University and University Hospital Schleswig-Holstein, Kiel, Germany; School of Medicine, University of Galway, Galway, Ireland; Department of Psychiatry and Psychotherapy, University Medicine Greifswald, Greifswald, Germany; Research Unit Analytical BioGeoChemistry, Helmholtz Munich, 85764 Neuherberg, Germany; CAU Innovation GmbH, Kiel University, Germany; Institute of Epidemiology, University Hospital Schleswig-Holstein, Campus Kiel, Kiel, Germany; Institute of Diabetes and Clinical Metabolic Research, University Hospital Schleswig-Holstein, Kiel, Germany; Inserm UMR-S 1256 nGERE (Nutrition-Genetics-Environmental Risks), Institute of Medical Research (Pôle BMS)-University of Lorraine, Vandœuvre-les-Nancy, France; German Center for Cardiovascular Diseases, Partner site North, Greifswald, Germany

## Abstract

Proton pump inhibitors (PPIs) alter the gut microbiome, but the contribution of these changes to systemic effects associated with PPI use remains unclear. Here, we show that PPI-associated microbiome remodelling induces systemic metabolic and transcriptional responses independently of direct drug exposure. Integrating human microbiome and metabolomics data with metabolic modelling, we identified consistent alterations in microbial metabolic functions and host–microbiome co-metabolism. Transfer of PPI-associated microbiota into germ-free mice reproduced key metabolic changes observed in humans, particularly in aromatic amino acid metabolism, and induced additional alterations in systemic lipid metabolism. These changes were accompanied by tissue-specific transcriptional responses in the colon, liver and brain, affecting metabolic and inflammatory pathways. Together, our findings identify transferable metabolic and transcriptional microbiome-mediated signatures that may contribute to the adverse systemic effects associated with PPI use.

## Introduction

Proton pump inhibitors (PPIs) are among the most widely prescribed acid-suppressive drugs and are a cornerstone treatment for gastroesophageal reflux disease, heartburn and peptic ulcers (1-3). Although highly effective, their prolonged use without a clear indication remains a clinical concern (4, 5). Observational studies have linked PPI use to enteric infections, chronic kidney disease and fractures (6, 7), type 2 diabetes (8, 9), inflammatory bowel disease (10) and neuropsychiatric disorders (11-13), although residual confounding and treatment of early symptoms before disease diagnosis complicate causal interpretation. A large randomized, placebo-controlled trial detected no significant increase in most assessed adverse outcomes over a median follow-up of three years, apart from a possible increase in enteric infections, although modest or longer-term risks could not be excluded (14). Gut microbiome alterations have been proposed as a potential link between PPI use and systemic outcomes (15, 16), motivating experimental investigation of whether PPI-associated microbiota can alter host metabolism and immune regulation independently of direct drug exposure.

By suppressing gastric acid secretion through inhibition of the gastric H⁺/K⁺-ATPase (17), PPIs alter the gastrointestinal environment and facilitate expansion of oral, facultative and oxygen-tolerant taxa in the lower gut (18, 19). Consistent with this, PPI users show altered microbial diversity and enrichment of upper gastrointestinal taxa including *Streptococcus*, *Enterococcus* and *Escherichia coli* (20). However, most studies of PPI-associated microbiome alterations have remained largely compositional, and it remains unclear how these taxonomic shifts translate into altered microbial metabolism or host physiology.

Constraint-based metabolic modelling provides a framework to connect microbiome composition to function by enabling prediction of metabolic fluxes and metabolite exchange within microbial communities (21). The availability of large-scale microbial reconstructions such as AGORA2 (22), together with community modelling platforms (23-25) and human whole-body metabolic reconstructions such as *Harvey* and *Harvetta* (26), now permits systems-level investigation of how microbiome-associated metabolic alterations may influence host physiology.

Here, we investigated whether PPI-associated microbiome alterations constitute a transferable functional state capable of shaping host metabolism and immune regulation. To address this, we integrated personalized microbiome and whole-body metabolic modelling across two human cohorts with serum metabolomics, *ex vivo* faecal cultivation and gnotobiotic mouse colonization. Across cohorts and experimental systems, PPI-associated microbiomes showed reproducible functional alterations that were more concordant than taxonomic changes and were characterized by a shift away from strictly anaerobic carbohydrate metabolism toward a more aerotolerant and amino acid-oriented metabolic state. These alterations persisted when transferring PPI-user microbiota into germ free mice, reshaping intestinal and systemic metabolite profiles and inducing tissue-specific host responses in recipient mice. Together, our findings provide evidence that PPI-induced changes in the gut microbiota can causally influence systemic host physiology.

## Results

### PPI use is associated with pronounced shifts in microbiome composition

To investigate functional differences associated with PPI use, we analysed two independent human datasets. The Kiel dataset combined 1,280 participants from the separately recruited PopGen and Food Chain Plus (FoCus) cohorts, with extensive phenotypic information and 16S rRNA gene-based microbiome profiles (27, 28). The MetaCardis dataset comprised 997 adults from Denmark, France and Germany across the cardiometabolic disease spectrum, including healthy controls and individuals with obesity, type 2 diabetes and ischemic heart disease (29). Extensive phenotypic information, shotgun metagenomic profiles, serum and urine metabolomics data, and detailed medication records including PPI exposure were available for MetaCardis.

We first examined whether PPI exposure was associated with shifts in microbiome taxonomic composition. Differential abundance analysis using MaAsLin2 for metagenomic data and ALDEx2 for 16S-based data revealed consistent directional associations across datasets after adjustment for demographic and clinical covariates (Fig. 1A,B). At the genus level, PPI-effect estimates were significantly correlated (Spearman ρ = 0.31, p = 0.037), with 73% of genera showing concordant directions of effect. Concordance also remained significant at the species level (Spearman ρ ≈ 0.17, p = 0.002), with approximately 75% of shared taxa exhibiting consistent directional effects. Two species, *Streptococcus salivarius* and *Streptococcus parasanguinis*, were significantly associated with PPI exposure across both cohorts (Fig. 1A,B).

**Figure 1.**
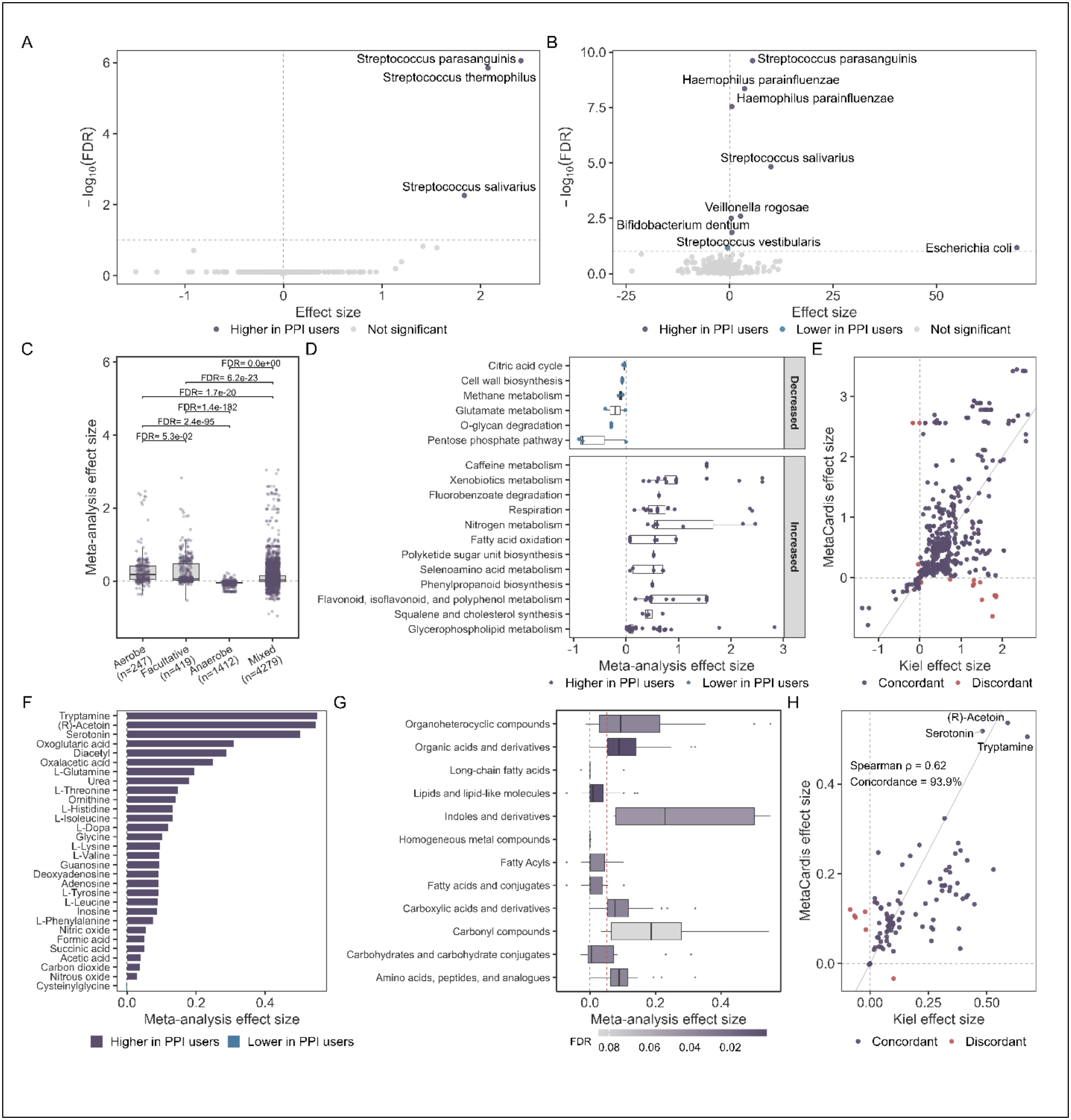
Cross-cohort microbiome and microbiome-derived functional signatures associated with proton pump inhibitor exposure. **(A)** Volcano plot of species-level associations with PPI use in the Kiel cohort. Each point represents a microbial species, plotted according to its ALDEx2 coefficient and −log10(FDR), with FDR calculated using the Benjamini–Hochberg method. Colours indicate significant positive or negative associations at FDR < 0.1. Dashed lines indicate zero coefficient and FDR = 0.1. **(B)** Volcano plot of species-level associations with PPI use in the MetaCardis cohort. Each point represents a microbial species, plotted according to its MaAsLin2 coefficient for PPI use and −log10(q-value). Colours indicate significant positive or negative associations at FDR < 0.1. Dashed lines indicate zero coefficient and FDR = 0.1. **(C)** Distribution of reaction-level meta-analysis effect sizes across microbial oxygen tolerance groups. Pairwise comparisons were assessed using two-sided Wilcoxon rank-sum tests with FDR correction. **(D)** Enriched metabolic subsystems among significantly altered reactions (FDR < 0.1). Only reactions significantly increased or decreased in proton pump inhibitor users are shown within the corresponding enrichment category. Boxplots show subsystem-level distributions of reaction meta-analysis effect sizes. **(E)** Cross-cohort comparison of significantly altered reaction abundance associations (meta-analysis FDR < 0.1). Points represent reactions plotted by cohort-specific effect sizes and coloured according to directional concordance between cohorts. Dashed grey lines indicate zero effect; diagonal lines indicate equal effect sizes across cohorts. **(F)** Top 30 significantly altered predicted metabolite secretion fluxes ranked by absolute meta-analysis effect size (FDR < 0.1). Colours indicate direction of association. **(G)** Enriched metabolite classes among predicted secretion fluxes associated with proton pump inhibitor exposure. Boxplots show distributions of meta-analysis effect sizes within each metabolite class; fill colour denotes FDR-adjusted enrichment significance. The dashed red line indicates the global median metabolite effect size. **(H)** Cross-cohort comparison of significantly altered predicted metabolite secretion fluxes (meta-analysis FDR < 0.1). Points represent metabolites plotted by cohort-specific effect sizes and coloured according to directional concordance between cohorts. Dashed grey lines indicate zero effect; diagonal lines indicate equal effect sizes across cohorts.

### Functional modelling reveals pronounced shifts in microbiome metabolic function in PPI users

Personalized constraint-based community models were constructed from microbiome composition using AGORA2 (22) to estimate microbiome functional potential. Meta-analysis identified 932 reaction-abundance associations with PPI exposure, of which 90.5% were positive and 9.5% negative, indicating an overall increase in model-inferred reaction abundance. Using AGORA2 strain-level oxygen-requirement annotations, reactions were classified according to whether their distribution across species models was preferentially associated with aerobes, facultative anaerobes or obligate anaerobes. These classes describe ecological associations and not the biochemical oxygen requirements of individual reactions. Aerobe- and facultative-associated reactions had higher PPI effect estimates than obligate-anaerobe-associated reactions, among significant reactions, all 58 aerobe-associated and 126 facultative-associated reactions were positive, whereas all 38 obligate-anaerobe-associated reactions were negative (Fig. 1C). This ordering was preserved across classification thresholds ranging from 0.10 to 0.30 and was consistent with the increased representation of aerotolerant oral taxa, including *Streptococcus salivarius*, in both cohorts. Reactions positively associated with PPI status were enriched for transport and exchange, glycerophospholipid, xenobiotic and respiration-related subsystems (Fig. 1D). The respiration annotation encompassed nitrate-, nitrite- and TMAO-reduction reactions, quinone-linked dehydrogenases and oxygen-dependent reactions, rather than exclusively aerobic respiration. AGORA2-annotated flavonoid and polyphenol subsystems were also enriched. Negatively associated reactions were enriched for O-glycan degradation and other host-derived carbohydrate-utilization pathways. Key reaction-level effects were directionally concordant between the Kiel and MetaCardis cohorts (Fig. 1E). Together, these results support reduced representation of functions preferentially encoded by obligate anaerobes alongside increased transport, respiratory and xenobiotic potential.

Analysis of predicted net secretion capacities identified 99 metabolites associated with PPI exposure, of which 93 had positive and six negative effect estimates. The largest positive effects involved aromatic amino-acid-related compounds, including tryptamine, serotonin, L-Dopa, indoleacetic acid and hydroxyphenyl derivatives, together with central-carbon and organic-acid metabolites such as (R)-acetoin, diacetyl, acetate, succinate, formate and lactate (Fig. 1F). At the metabolite-class level, indoles, carbonyl compounds and other organoheterocyclic or amino-acid-related metabolites showed the largest positive effect-size distributions, whereas carbohydrate- and fatty-acid-related classes showed smaller effects (Fig. 1G). Secretion-capacity effects were concordant between cohorts (Spearman ρ = 0.62; 93.9% directional concordance; Fig. 1H).

### Functional alterations in PPI-associated microbiome metabolism impact host pathways

To assess how PPI-associated microbiomes could alter host–microbiome co-metabolism, personalized microbial community reconstructions were integrated with whole-body metabolic models, and maximal demand fluxes were calculated for metabolites in the blood compartment. These predicted blood-compartment fluxes showed substantial agreement between cohorts (Spearman ρ = 0.49), with 71.6% of metabolites showing concordant effect directions (Fig. 2A). Significant meta-analysis associations were predominantly positive and concentrated in amino acid-related pathways, including aromatic amino acid, histidine and imidazole, branched-chain amino acid and lysine metabolism (Fig. 2B). This pattern was consistent with the increased amino acid-catabolic and proteolytic potential inferred from the microbial community models.

**Figure 2.**
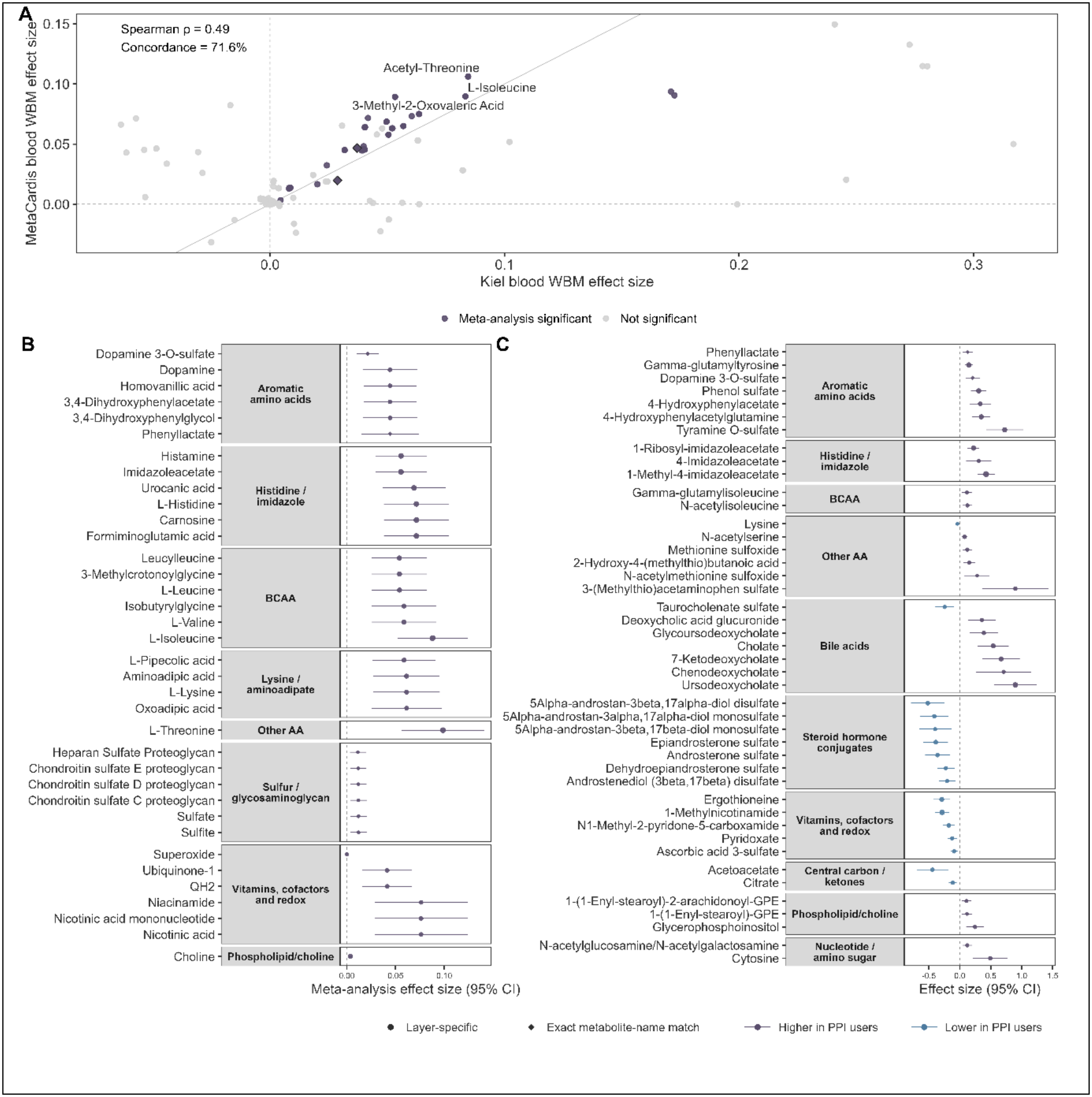
Whole-body modelling across cohorts and serum metabolomics identify overlapping PPI-associated metabolic themes. **(A)** Meta-analysis of reaction-level blood-flux associations from personalized whole-body metabolic models in the Kiel and MetaCardis cohorts. Each point represents a reaction plotted according to its cohort-specific PPI effect estimate. Purple points indicate robust meta-analysis signals (FDR < 0.1, concordant effect direction between cohorts, Cochran’s Q *P* ≥ 0.05 and I² ≤ 25%), whereas grey points indicate other tested reactions. Diamonds denote robust reactions with an exact metabolite-name match in the measured serum metabolomics data. Dashed lines indicate zero effect, and the diagonal line indicates equal effect sizes between cohorts. Spearman’s ρ and the fraction of reactions with concordant effect directions are shown. The three top-ranked robust signals are labelled. **(B)** Representative blood-flux associations identified by fixed-effect meta-analysis of the Kiel and MetaCardis whole-body models, grouped by curated metabolic theme. Near-identical model estimates were collapsed, with one representative metabolite displayed for each coupled flux signature. **(C)** Representative serum metabolites significantly associated with PPI exposure in the MetaCardis cohort, grouped according to the same metabolic themes. In **B** and **C**, points and horizontal lines show effect estimates and 95% confidence intervals. Purple and blue indicate higher and lower predicted blood flux or serum abundance, respectively, in PPI users. Circles denote layer-specific signals, whereas diamonds indicate exact metabolite-name matches between the modelling and measured-metabolomics layers. Point size is proportional to −log₁₀(FDR). Direct medication measurements and unidentified features were excluded from the displayed serum-metabolomics results.

In addition to predicting microbial metabolite secretion, we analysed serum metabolomics data from the MetaCardis cohort to investigate the potential impact of PPI-associated microbiome alterations on the host serum metabolome. Aromatic amino acid metabolism represented the strongest point of overlap across the microbial, whole-body modelling and measured-metabolomics layers. Personalized microbiome models predicted increased secretion of tryptamine, serotonin, L-Dopa, indole-related compounds and hydroxyphenyl derivatives. Consistent with these predictions, whole-body models indicated increased maximal blood-compartment demand capacity of aromatic amino acid products, including dopamine 3-O-sulfate and phenyllactate, while serum metabolomics showed higher concentrations of phenol sulfate, tyramine O-sulfate, dopamine 3-O-sulfate, 4-hydroxyphenylacetate, 4-hydroxyphenylacetylglutamine and phenyllactate in PPI users (Fig. 2B,C). Dopamine 3-O-sulfate and phenyllactate were exact metabolite-name matches between the whole-body modelling and serum-metabolomics layers. Together with the aromatic metabolites identified in the community models, these findings indicate coordinated changes in microbial aromatic amino acid transformation and subsequent host conjugation and systemic handling reflected in serum metabolomics.

Additional overlap was observed for histidine and branched-chain amino acid metabolism, with increased predicted maximal blood-compartment demand capacity of histamine, imidazole derivatives, leucine, isoleucine and valine, accompanied by corresponding serum associations (Fig. 2B,C). Serum metabolomics further revealed increased concentrations of primary and secondary bile acids, including cholate, ursodeoxycholate, chenodeoxycholate and glycoursodeoxycholate, consistent with altered microbial transformation and enterohepatic processing. In contrast, several androgen-derived steroid sulfates and metabolites related to central carbon, NAD⁺, vitamin and redox metabolism were decreased. Although microbial secretion fluxes, whole-body blood fluxes and circulating metabolite concentrations represent distinct quantities, their convergence across metabolic pathways supports a systemic PPI-associated signature centred on aromatic amino acid metabolism and enhanced host processing of microbiome-derived metabolites. Full taxonomic, microbiome-modelling, serum-metabolomics and whole-body-model association results are provided in Supplementary Data 1.

### PPI-associated microbiomes exhibit transferable metabolic alterations in vitro and in vivo

Having observed pronounced model-predicted shifts in the metabolic output of PPI-associated microbiomes and corresponding alterations in the serum metabolome of the host, we next performed functional characterization of these microbiomes *in vitro* and *in vivo* to distinguish microbiota-mediated effects from host-related differences or direct effects of PPIs on host metabolism. Faecal samples from three pantoprazole users and three controls matched by sex and approximate year of birth were used for ex vivo incubation and colonization of germ-free mice. Each group included two men and one woman; donor characteristics and recorded diagnoses are provided in Table 1 (in Methods). For *ex vivo* experiments, faecal material was cultured for 96 h at 37 °C in pre-reduced mGAM medium under strictly anaerobic conditions, with daily collection of bacterial pellets for 16S rRNA gene sequencing and supernatants for metabolomics. To assess host effects, faecal material from the same donors was transplanted into germ-free mice. Two weeks post colonization under gnotobiotic conditions, faecal and caecal contents were collected for 16S rRNA gene sequencing and non-targeted metabolomics, while colon, liver and brain tissues were retained for transcriptomic analysis. Microbiome profiling demonstrated that donor identity remained a major determinant of community composition in both *ex vivo* cultures and recipient mice. Samples derived from the same donor clustered according to donor origin in ordination space (Fig. 3A,B) and exhibited lower Bray– Curtis dissimilarities than samples originating from different donors (Fig. 3C,D), indicating that donor-specific microbiome signatures were maintained following *ex vivo* cultivation and gnotobiotic transfer.

**Figure 3.**
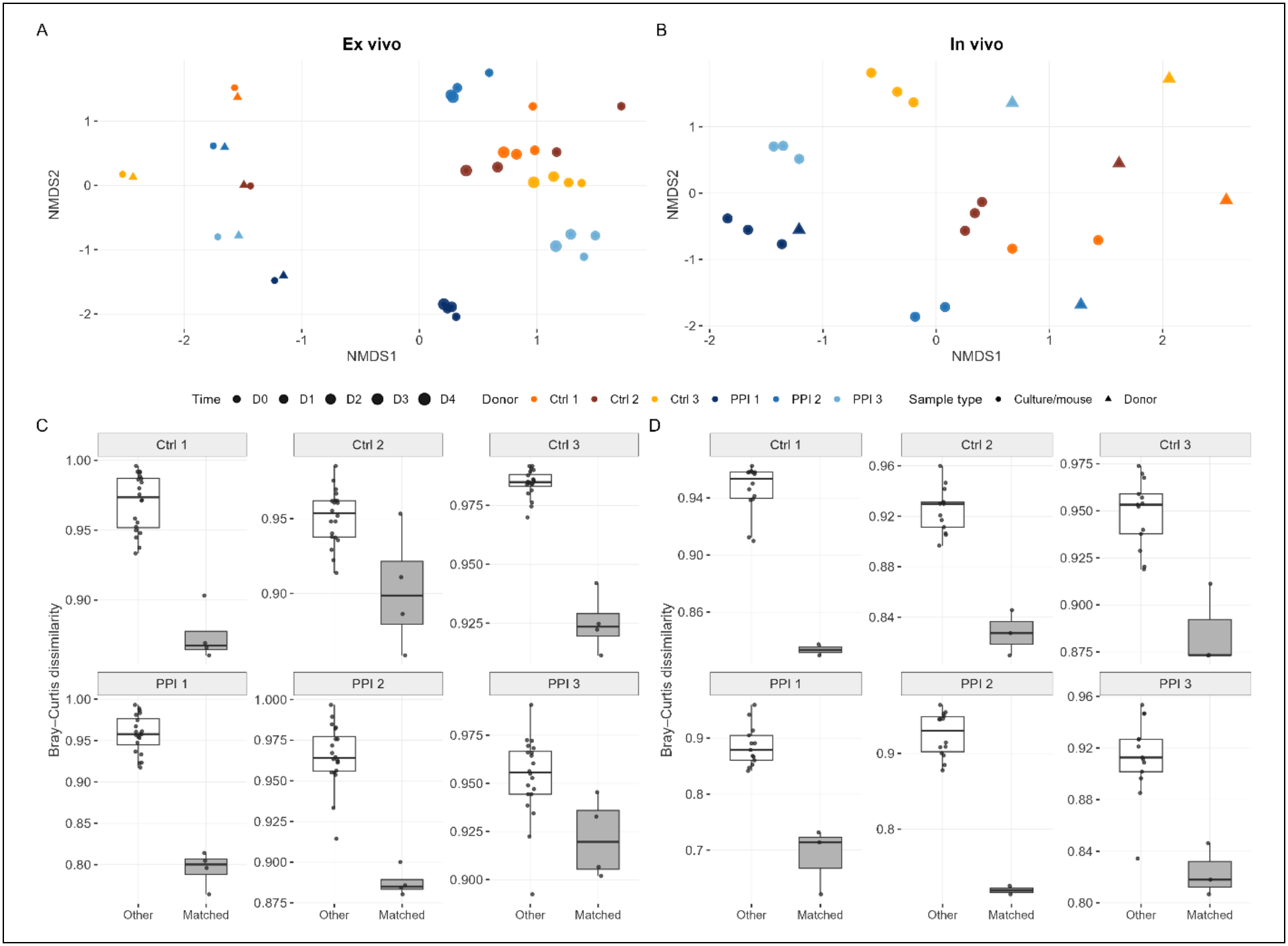
Donor-specific microbiome similarity after ex vivo cultivation and transfer to germ-free mice. **(A,B)** Non-metric multidimensional scaling (NMDS) of Bray–Curtis dissimilarities for ex vivo cultures (A) and mouse recipients (B). Colours identify the six donors; triangles denote original donor samples. In A, point size indicates culture day. **(C,D)** Bray–Curtis dissimilarities between each original donor sample and cultures (C) or mouse recipients (D) originating from the same donor (“Matched”) or other donors (“Other”). Each point represents one donor-to-sample comparison. Matched samples were generally more similar to their donor of origin.

**Table 1.** Characteristics of faecal donors used for ex vivo cultivation and germ-free mouse colonization. Three pantoprazole users and three controls were matched by sex and year of birth (within two years). Recorded diagnoses do not necessarily indicate the reason for PPI treatment. M, male; F, female.

| **Matched pair** | **Group** | **Birth year** | **Sex** | **Recorded diagnoses** |
|---|---|---|---|---|
| A | Pantoprazole | 1958 | M | Varicose veins; irritable bowel syndrome |
| A | Control | 1958 | M | No diagnoses recorded |
| B | Pantoprazole | 1965 | F | Urticaria; psoriasis; rheumatoid arthritis |
| B | Control | 1964 | F | Flat feet; nephrolithiasis (2017) |
| C | Pantoprazole | 1995 | M | Allergic rhinitis; psoriasis; periodontitis |
| C | Control | 1993 | M | Haemorrhoids |

### PPI-associated microbiota induce intestinal and systemic metabolic alterations consistent with changes observed in PPI users

To determine whether PPI-associated microbiota retained distinct metabolic properties outside the donor host, we profiled *ex vivo* culture supernatants as well as intestinal contents and serum from recipient mice using complementary NMR- and LC–MS-based approaches (Fig. 4; Supplementary Data 3). Global metabolomic variation was explored using PCA across cultures, intestinal compartments and serum (Fig. 4A–E). After each culture was centred to its day-0 profile, temporal changes measured by NMR and HILIC LC–MS were strongly concordant across matched samples (Procrustes r = 0.857, permutation P = 10⁻⁴).

**Figure 4.**
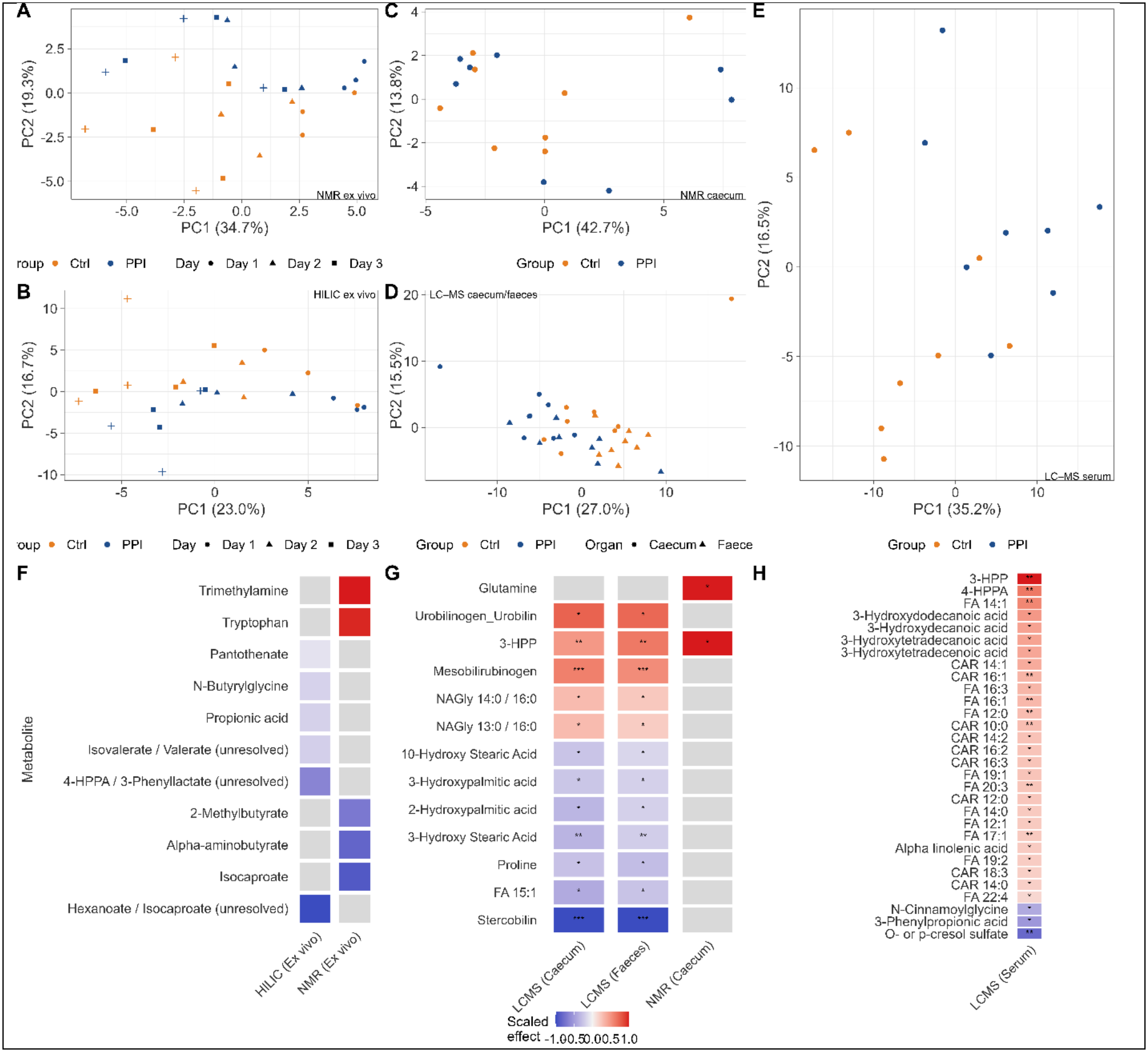
Multimodal metabolomic profiling identifies group-associated metabolic variation across *ex vivo* cultures, intestinal compartments and serum. **(A–E)** Principal component analysis (PCA) of metabolomic profiles from *ex vivo* cultures measured by NMR **(A)** and HILIC LC– MS **(B)**, caecal samples measured by NMR (C), combined caecal and faecal samples measured by RP LC–MS **(D)**, and serum measured by RP LC–MS **(E)**. Points are coloured by experimental group (Ctrl or PPI). In A and B, symbols indicate the sampling day; in D, symbols distinguish caecal and faecal samples. The percentages of variance explained by the first two principal components are shown on the axes. **(F–H)** Heatmaps of group-associated metabolites. Colours represent effect estimates standardized within each dataset to allow comparison across platforms with different effect scales; red indicates higher abundance in the PPI group and blue indicates lower abundance. Metabolites are ordered from the highest to the lowest standardized effect. Asterisks indicate significance thresholds (*FDR < 0.10, **FDR < 0.05, ***FDR < 0.01), while grey cells indicate that the metabolite did not meet the corresponding inclusion criterion in that dataset. Metabolite names were harmonized across platforms, including the use of 3-HPP and 4-HPPA. **(F)** Ex vivo metabolites measured by HILIC LC–MS and NMR that were nominally associated with group (P<0.05). No feature remained significant at FDR < 0.1. **(G)** Intestinal metabolites measured by RP LC–MS in caecum and faeces and by NMR in caecum. For the RP LC–MS datasets, significance reflects the overall group effect across intestinal compartments. (H) The 30 serum RP LC–MS metabolites with FDR < 0.10 showing the largest absolute group effects.

At the individual-metabolite level, *ex vivo* NMR showed nominally higher tryptophan and trimethylamine and lower alpha-aminobutyrate, 2-methylbutyrate and isocaproate in PPI-associated cultures (Fig. 4F). None remained significant after multiple-testing correction, and no HILIC metabolite reached FDR < 0.1. Recipient-mouse intestinal metabolomics revealed clearer individual changes. Combined caecal and faecal RP LC–MS identified 12 metabolites at FDR < 0.1 (Fig. 4G). Stercobilin was markedly depleted, whereas mesobilirubinogen and the urobilinogen/urobilin feature were increased, consistent with redistribution among intestinal bilirubin products. 3-(3-Hydroxyphenyl)propionic acid (3-HPP) increased 6.9-fold by RP LC–MS (FDR = 0.023), with independent support from caecal NMR (score difference = 1.53, FDR = 0.090). Caecal NMR additionally showed higher glutamine, while intestinal RP LC–MS identified lower hydroxylated fatty acids and higher selected N-acylglycine species (Fig. 4G). The 3-HPP result extended the broader aromatic-metabolism signal observed in human serum and predicted by the microbial models. Serum RP LC–MS identified 30 metabolites at FDR < 0.1 (Fig. 4H). 3-HPP and 3-(4-hydroxyphenyl)propionic acid (4-HPPA) increased 6.9-fold and 3.4-fold, respectively, extending the aromatic signal into the circulation. In contrast, 3-phenylpropionic acid, cresol sulfate and N-cinnamoylglycine were reduced, indicating redistribution among related aromatic metabolites. Most other significant serum changes were increases in medium- and long-chain fatty acids, hydroxylated fatty acids and acylcarnitines.

### PPI-associated microbiota induce coordinated immune and metabolic host responses

To determine whether transfer of PPI-associated microbiota was associated with host transcriptional responses, we analysed gene expression of the mice in liver, colon and brain. Global transcriptional variation was visualized by PCA (Fig. 5A–C). DESeq2 identified 650 differentially expressed genes in liver, compared with three in colon and four in brain (FDR < 0.1; Fig. 5D–F; Supplementary Data 4), indicating that the strongest gene-level response occurred in liver.

**Figure 5.**
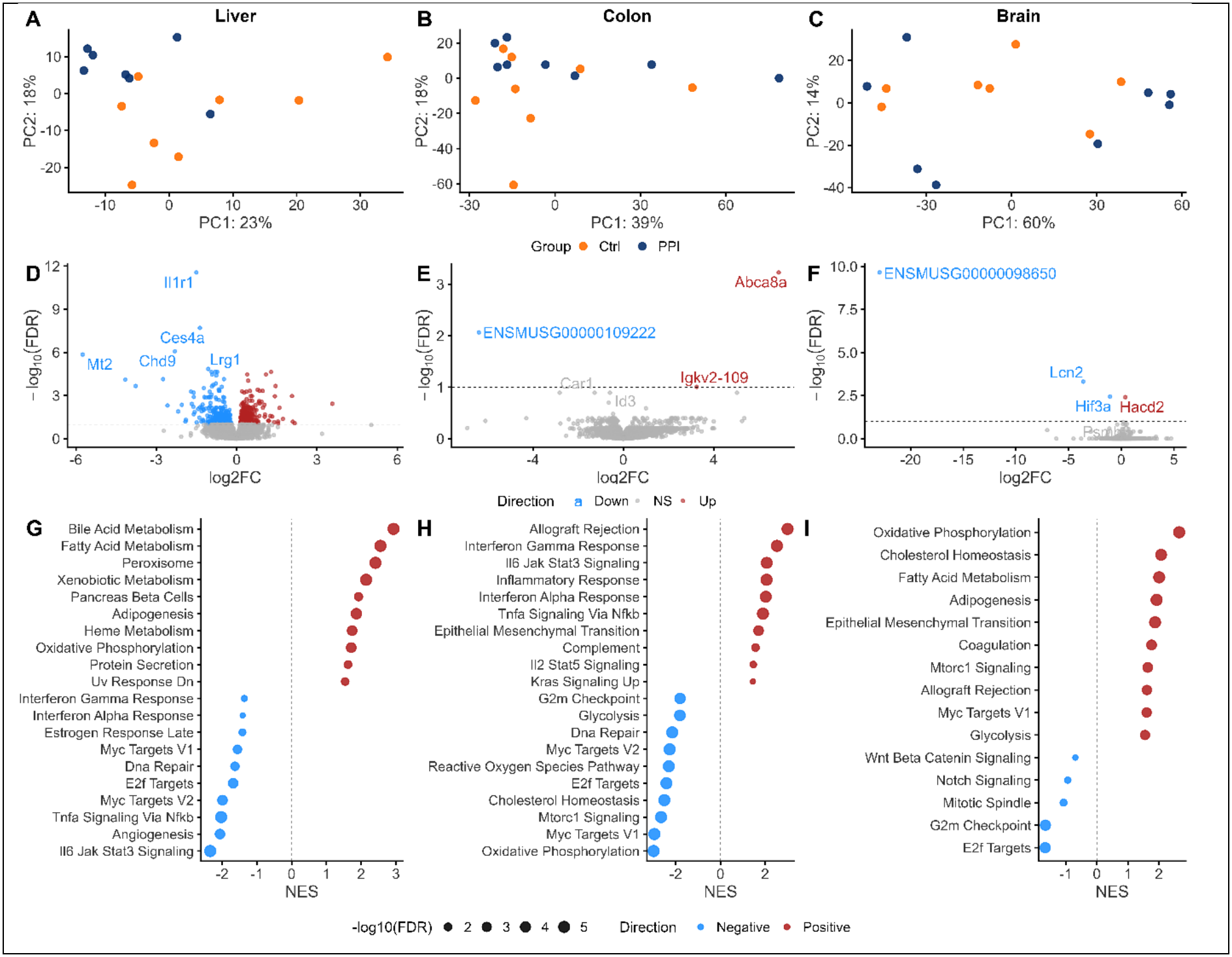
Organ-specific transcriptional responses to PPI-associated microbiota. **(A–C)** PCA of variance-stabilized RNA-seq counts from liver **(A)**, colon **(B)** and brain **(C)**, coloured by microbiota group. **(D–F)** Volcano plots of differential expression between recipients of PPI-associated and control microbiota in liver **(D)**, colon **(E)** and brain **(F)**. Positive log2 fold changes indicate higher expression in PPI recipients. The dashed horizontal line indicates FDR = 0.1; colours denote significantly increased, significantly decreased or non-significant genes. **(G–I)** Hallmark GSEA results for liver **(G)**, colon **(H)** and brain **(I)**. Dot position indicates the normalized enrichment score (NES), dot size indicates −log10(FDR), and colour indicates enrichment direction. Positive NES values denote enrichment among genes with higher expression in PPI recipients, whereas negative values denote enrichment among genes with lower expression.

In liver, GSEA showed positive enrichment of bile acid, fatty acid, peroxisomal, xenobiotic, haem-metabolism and oxidative-phosphorylation programmes. Inflammatory pathways, including TNFα–NFκB, IL6–JAK–STAT3 and interferon responses, were negatively enriched, together with MYC, E2F and DNA-repair programmes (Fig. 5G). Gene Ontology analysis supported these patterns: genes increased in PPI recipients were enriched for small-molecule, fatty-acid, lipid and amino-acid catabolism, whereas decreased genes were enriched for acute-phase, complement, humoral immune, Toll-like receptor and TNF–NFκB processes. Iron-ion homeostasis terms were also enriched among decreased genes (Supp. Fig. 1A). In colon, inflammatory, interferon, IL6–JAK–STAT3, TNFα–NFκB and complement pathways were positively enriched. Oxidative phosphorylation, cholesterol homeostasis and mTORC1 signalling were negatively enriched, together with cell-cycle and proliferative programmes including E2F, G2M, MYC and DNA repair (Fig. 5H). At gene level, *Abca8a* and *Igkv2-10S* were increased in PPI recipients (Supp. Fig. 1B). *Abca8a*, an ATP-binding cassette transporter associated with lipid handling, was largely undetectable in control recipients but expressed in most PPI recipients, suggesting a potential link between altered microbial metabolism and host lipid or bile-acid handling. Brain responses were weaker. Metabolic pathways, including oxidative phosphorylation, cholesterol homeostasis and fatty-acid metabolism, were positively enriched, whereas E2F, G2M, mitotic-spindle and Wnt/Notch programmes were negatively enriched (Fig. 5I). Among the four differentially expressed genes in brain, *Hif3a*, *Lcn2* and *Commd1b* were decreased, whereas *Hacd2* was increased in PPI user stool recipients (Fig. 5F; Supp. Fig. 1C,D). *Lcn2* was also decreased in liver, representing a shared gene-level response between the two tissues (Supp. Fig. 1D). Unlike colon, brain did not show a coordinated interferon, TNF or IL6 signature. Thus, although the overall responses were organ-specific, negative enrichment of cell-cycle and proliferative programmes recurred across tissues. Reactome analyses supported the same broad tissue-specific patterns (Supplementary Data 4).

### Microbiome-associated metabolic pathways are linked to tissue-specific host transcriptional responses

To relate microbial community structure to metabolites and host transcription, we performed two linked analyses. We first tested genus–metabolite associations in matched experimental samples, accounting for treatment group and repeated samples from each donor. We then tested whether metabolites altered in recipient mice were associated with tissue gene expression after adjustment for recipient group and summarized the gene-level associations by pathway enrichment (Methods). Across *ex vivo* and mouse metabolomics datasets, 190 genus–metabolite pairs met FDR < 0.1, most frequently involving *Colidextribacter* (99 associations), *Alistipes* (22) and *Agathobaculum* (19). In leave-one-donor-out refits, 189 of 190 associations retained their direction after each donor was excluded (Supplementary Data 5). Aromatic metabolites provided a cross-platform signal. Higher caecal 3-HPP, which was increased in PPI recipients, was associated with higher *Alistipes* abundance (Figs. 4G and 6A). Gene-level association statistics for 3-HPP were concordant between serum and caecal measurements in colon and liver (Spearman ρ = 0.75–0.96; Fig. 6B). In colon, higher caecal 3-HPP was associated with negative enrichment of E2F targets, G2M checkpoint, MYC targets, mitotic spindle, protein secretion and mTORC1 gene sets (Fig. 6C). In other words, genes in these sets tended to have lower expression at higher 3-HPP abundance after adjustment for recipient group. Other aromatic metabolites showed distinct, sometimes opposing pathway associations (Fig. 6C,D). *Alistipes* and *Colidextribacter* were also inversely associated with caecal aromatic amino acids (Fig. 6A). *Colidextribacter* was associated with lipid metabolites, including serum lysophospholipids and fatty acids (Fig. 6A; Supplementary Data 5). Higher serum FA 22:4 was associated with negative enrichment of colonic cholesterol homeostasis and biosynthesis, E2F, G2M, MYC and mTORC1 gene sets (Fig. 6C). *Agathobaculum* was positively associated with intestinal bile acid derivatives and serum DCA (Fig. 6A). Bilirubin products showed differing transcriptional associations: mesobilirubinogen and urobilinogen/urobilin had broadly similar pathway patterns in liver, whereas stercobilin frequently showed the opposite direction (Fig. 6C,D). In colon, higher stercobilin was associated with negative enrichment of interferon, TNF–NFκB, inflammatory-response and IL6–JAK–STAT3 gene sets, and positive enrichment of oxidative phosphorylation (Fig. 6C). It is important to note that these analyses identify associations and cannot establish that a taxon produces a metabolite or that a metabolite causes the transcriptional response. Together, these analyses link variation in single gut bacterial genera to metabolite profiles and highlight 3-HPP as a metabolite associated with both *Alistipes* abundance and transcriptional programs in the colon and liver.

**Figure 6.**
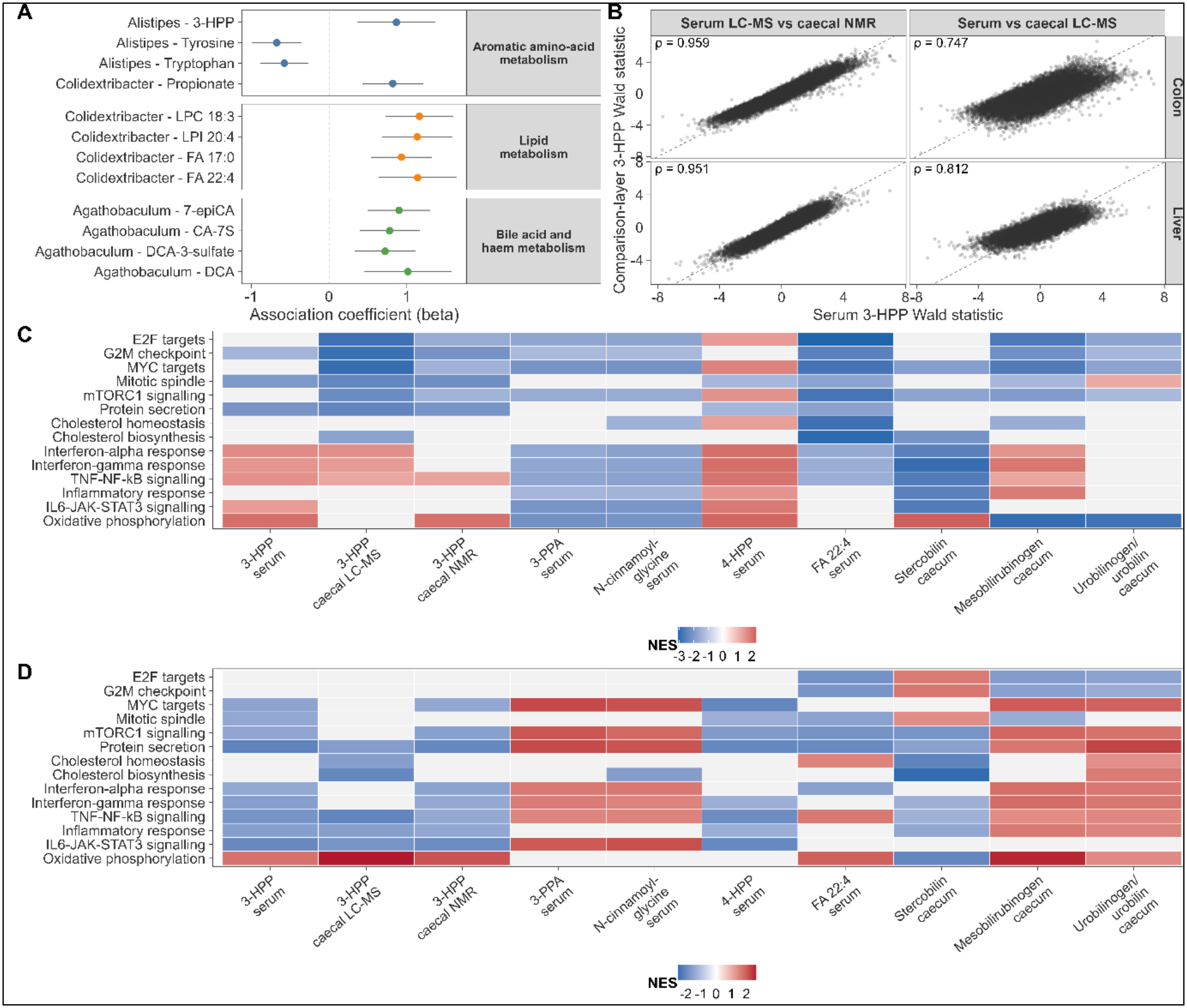
Cross-layer associations between microbial genera, metabolites and host transcriptional programmes. **(A)** Selected genus–metabolite associations grouped into aromatic amino-acid, lipid, and bile acid or haem-related metabolic themes. Points indicate adjusted rank-standardized association coefficients (β), and horizontal lines show 95% confidence intervals. **(B)** Cross-platform concordance of gene-level DESeq2 Wald statistics for 3-(3-Hydroxyphenyl)propionic acid (3-HPP) in colon and liver. Serum LC–MS-derived statistics were compared with those obtained using caecal NMR or caecal LC–MS measurements. Each point represents one gene, dashed lines indicate the identity line, and ρ denotes Spearman’s rank correlation coefficient. **(C–D)** Associations between metabolite abundance and transcriptional pathways in colon **(C)** and liver **(D)**, assessed by DESeq2-derived gene set enrichment analysis. Tile colour represents the normalized enrichment score (NES), with red indicating positive and blue indicating negative pathway associations with metabolite abundance. Coloured cells indicate associations with family-level FDR < 0.10; grey cells indicate that no significant association was retained for the corresponding metabolite–pathway combination.

**Figure 7.**
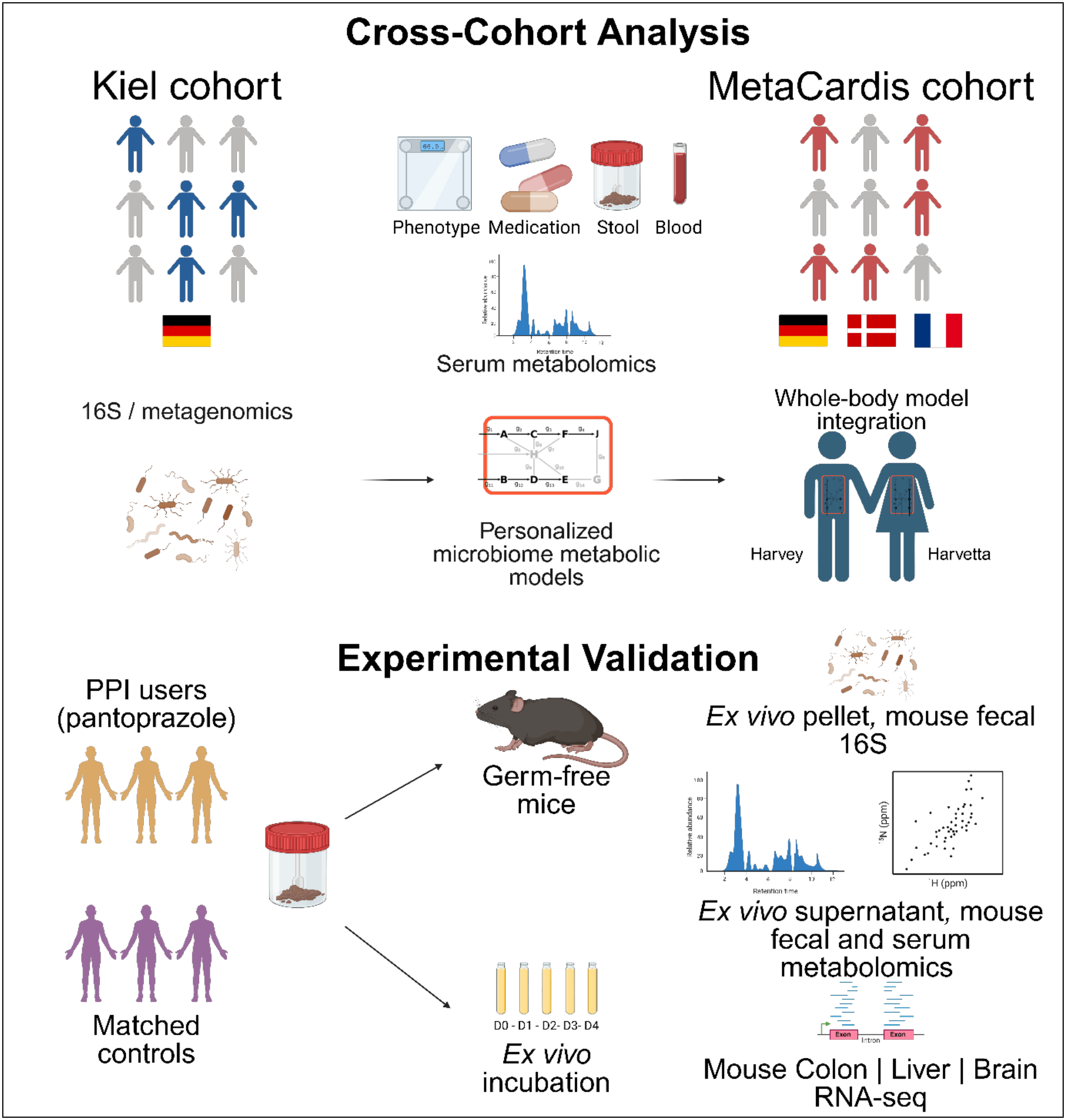
Study design and analytical workflow.

## Discussion

In this study, we identified a reproducible functional remodelling of the gut microbiome in PPI users and show that key aspects of this phenotype are transferable with the microbiota. Across two independent human cohorts, PPI use was associated with consistent shifts in model-predicted microbiome metabolism accompanied by changes in the serum metabolome. While enrichment of oral and facultatively anaerobic taxa, including *Streptococcus*, *Haemophilus* and *Escherichia coli*, is consistent with previous studies (20), our data suggest that the consequences of PPI exposure are captured more robustly at the level of microbial metabolic function than by individual taxa. Altered functions converged particularly on aromatic amino acid metabolism, fermentation-associated pathways, microbial bilirubin reduction and host metabolism of microbiome-derived compounds. Importantly, transplantation of PPI-associated microbiota into germ-free mice recapitulated selected metabolic features and induced organ-specific host transcriptional responses without direct PPI exposure, supporting a microbiota-mediated contribution to systemic effects of PPIs. On a functional level, the PPI-associated microbiomes showed increased overall metabolic activity alongside a broader ecological shift from saccharolytic and strictly anaerobic functions towards increased amino acid-catabolic and aerotolerant metabolic potential. This was supported by reaction-level analyses, predicted secretion profiles and serum metabolomics. These functional changes extend previous observations of increased oral and facultatively anaerobic taxa and may result from facilitated passage of oral bacteria into the colon, direct effects of PPIs on bacterial metabolism, or both (30). The resulting pattern resembles metabolic features associated with impaired host health, including enhanced amino acid catabolism and reduced anaerobic function (31, 32).

Aromatic amino acid metabolism provided one of the strongest links between microbiome modelling, human serum metabolomics and gnotobiotic experiments. Gut microbial metabolism of tryptophan, tyrosine and phenylalanine generates bioactive compounds that can influence intestinal barrier function, immunity and systemic metabolism (33-36). In PPI users, microbiome models predicted increased secretion potential for aromatic amino acid-derived metabolites, while serum metabolomics showed higher concentrations of several host-conjugated phenolic and indolic compounds, including indoxyl sulfate, phenol sulfate and hydroxyphenyl derivatives. Similar associations between PPI use and indoxyl sulfate or phenylacetylglutamine have been reported in patients with chronic kidney disease (37), while experimental evidence suggests that PPIs may also increase indoxyl-sulfate exposure by altering host conversion pathways (38). Additionally, several aromatic co-metabolites have been linked to cardiovascular disease and other adverse outcomes (39-41). Transfer of PPI-associated microbiota from human donors into germ-free mice reproduced this altered aromatic metabolic output. In particular, 3-(3-hydroxyphenyl)propionic acid (3-HPP) increased in intestinal contents and serum and was detected by complementary metabolomics approaches. Although 3-HPP can derive from several aromatic substrates and therefore cannot be assigned to a single precursor pathway (33, 42), its accumulation is consistent with altered microbiota-dependent processing or host handling. Importantly, the phenotype did not represent a general increase in aromatic degradation: hydroxyphenylpropionate-type metabolites increased, whereas 3-phenylpropionic acid and cresol sulfate decreased. However, reduced cresol sulfate also agrees with the loss of strictly anaerobe-associated functions, as microbial p-cresol formation in anaerobic Clostridia involves an oxygen-sensitive glycyl-radical enzyme (43, 44). Cross-layer integration further linked *Alistipes* to 3-HPP, which was associated with reduced proliferative and biosynthetic programmes in colon, supporting a connection between microbial aromatic metabolism and tissue-specific host responses. Higher 3-HPP was associated with lower hepatic inflammatory gene-set expression (Fig. 6D), consistent with anti-inflammatory effects reported in literature, although these associations did not establish a specific 3-HPP mediated response in the liver (45, 46).

Another phenotype consistent across human cohort data and gnotobiotic experiments involved microbial metabolism of the haem-catabolite bilirubin. Following hepatic conjugation and biliary excretion, bilirubin undergoes extensive microbial transformation in the intestine (47-49). The identification of microbial bilirubin reductase BilR established bilirubin-to-urobilinogen conversion as a defined microbial function, whereas terminal steps towards stercobilin remain incompletely resolved (47). In PPI user stool-recipient mice, mesobilirubinogen and urobilinogen/urobilin increased whereas stercobilin was strongly depleted, indicating redistribution within microbial bilirubin catabolism rather than uniform pathway suppression. Similar depletion of terminal bilirubin metabolites has recently been reported in IBD and associated with depleted *Lachnospiraceae*, including *Blautia* (50), different *Blautia* species also tended to be reduced in PPI users in both human cohorts. As microbial bilirubin metabolism can affect intestinal barrier and immune functions (49, 51), these findings identify altered exposure to haem-derived microbial metabolites as another potential interface between PPI-associated dysbiosis and host physiology. Bilirubin catabolites also showed coherent host transcriptional associations, particularly in colon, linking this metabolic phenotype to inflammatory and oxidative-phosphorylation programmes. In liver and brain, bilirubin catabolites were also associated with iron-transport programmes, with *Lcn2* among the leading-edge genes. LCN2 is a siderophore-binding acute-phase protein that sequesters iron and contributes to iron homeostasis during inflammation (52, 53).

Beyond aromatic amino acid and bilirubin metabolism, recipient-mouse serum showed coordinated increases in medium- and long-chain acylcarnitines, several of which were associated with *Colidextribacter*, consistent with altered mitochondrial fatty-acid handling or substrate utilization (54, 55). Such acylcarnitine accumulation can arise when fatty-acid supply exceeds oxidative capacity or when β-oxidation becomes incompletely coupled to downstream mitochondrial metabolism (54), and may therefore reflect altered mitochondrial substrate flux. At the transcriptional level, oxidative-phosphorylation pathways were negatively enriched in colon but positively enriched in liver and brain, whereas inflammatory programmes increased in colon but decreased in liver. These opposing transcriptional patterns are inconsistent with a uniform systemic response and instead suggest organ-specific differences in metabolic and inflammatory regulation. At gene level, increased colonic *Igkv2-10S* was consistent with altered mucosal antibody expression, while brain changes in *Hacd2*, *Hif3a* and *Lcn2* involved genes previously linked to mitochondrial disease or cognitive-disorganization traits, anxiety-related behaviour and stress-induced anxiety, respectively (56-59). These associations provide biological context, but behavioural outcomes were not assessed in the recipient mice.

Bile-acid metabolism showed a related but weaker signal across datasets. In recipient mice, *Agathobaculum* was associated with several intestinal bile-acid derivatives and circulating DCA. In MetaCardis, multiple primary and secondary bile acids were associated with PPI use, consistent with previous reports (60). Recipient mice also showed directional changes in multiple bile acids, although individual species did not remain significant after multiple-testing correction and directions were not consistently conserved between humans and mice. This is not unexpected given substantial interspecies differences in bile-acid composition (61). Notably, *Abca8a* showed non-zero colonic expression in seven of eight PPI recipients compared with one of eight controls and has been implicated in sterol trafficking and cholesterol and taurocholate efflux (62). Together with the bilirubin phenotype, these findings suggest that PPI-associated microbiota alter microbial and host processing of hepatobiliary metabolites. Given the roles of bile acids in glucose and lipid metabolism, epithelial barrier function and mucosal immunity (63, 64), such changes provide plausible links to metabolic and inflammatory effects associated with PPI use.

The transplantation experiment provides evidence that PPI-associated donor microbiota can induce metabolic and transcriptional host responses without direct drug exposure, while the *ex vivo* experiments suggest that these microbiomes retain metabolic differences outside the host. Several limitations should nevertheless be considered. The human analyses were observational, metabolic modelling estimated functional potential rather than realised *in vivo* flux, and serum metabolomics reflects a mix of microbial production together with host absorption, transformation and clearance. Transplantation included only three PPI and three control donor microbiomes, with recipient mice nested within donors, limiting generalisability, while the short colonisation period may not capture longer-term adaptation. Humanised mouse models can reproduce donor-specific metabolic features but remain subject to cross-species engraftment biases (65, 66). Studies with more donors, longer colonisation and targeted perturbation of candidate taxa or metabolites will be required to resolve the mechanisms underlying the transferred phenotype.

In summary, our results support a model in which PPI use reshapes gut microbiome composition and metabolic function, with consequences extending to host metabolism and gene regulation. The PPI-associated microbiome displayed a coherent metabolic phenotype across cohorts centred on altered aromatic amino acid co-metabolism and hepatobiliary processing. Transfer into gnotobiotic mice reproduced key metabolic features and induced tissue-specific host responses without direct PPI exposure, supporting microbiome remodelling as a potential contributor to metabolic and inflammatory responses associated with PPI use. These findings suggest that alterations in the gut microbiome may mediate part of the adverse health outcomes associated with long-term PPI use, providing a potential mechanistic link between PPI exposure and its reported systemic effects. To our knowledge, this is the first experimental demonstration that transfer of PPI-conditioned human gut microbiota can drive metabolic shifts in gnotobiotic mice. These findings underscore the value of examining microbial metabolic function alongside community composition and provide a basis for testing whether similar shifts contribute to the potential adverse outcomes reported in long-term PPI users.

## Data availability statement

Raw 16S rRNA gene amplicon sequencing data from donor faeces, ex vivo microbiota cultures and recipient mice, together with RNA-seq reads from mouse brain, colon and liver, have been deposited in the European Nucleotide Archive under accession PRJEB129040 (https://www.ebi.ac.uk/ena/browser/view/PRJEB129040). ASV count tables, taxonomic annotations, abundance tables, sample metadata and Bray–Curtis source data are provided in Supplementary Data 2. Raw RNA-seq reads have been deposited in the European Nucleotide Archive under project accession PRJEB129040. The processed gene-level count matrix and associated experimental metadata have been deposited in the ArrayExpress collection in BioStudies under accession E-MTAB-17878.

LC–MS/MS metabolomics data have been deposited in the NIH Common Fund’s National Metabolomics Data Repository, Metabolomics Workbench (67), under project PR003489. The project comprises studies ST005391 (ex vivo cultures), ST005392 (caecal and faecal contents) and ST005393 (serum samples), accessible through https://doi.org/10.21228/M8VT1GP9D. Processed NMR intensity tables and sample metadata for mouse caecal contents and ex vivo culture supernatants are provided in Supplementary data 6, with acquisition and processing protocols described in Methods. Numerical data underlying the figures are provided in the accompanying Source Data files.

Samples and data from the PopGen cohort were provided by the PopGen Biobank (Schleswig-Holstein, Germany). Access is available through a structured application procedure via the PopGen 2.0 Network (https://portal.popgen.de/), subject to approval and applicable data-use conditions. Publicly released MetaCardis processed omics data, associated metadata and medication profiles are available in Supplementary Tables 9–14 accompanying Fromentin et al. (2022; https://doi.org/10.1038/s41591-022-01688-4) and through the MetaDrugs repository on Zenodo (https://doi.org/10.5281/zenodo.6242715). Access to additional, non-public participant-level phenotypic data may be requested from the MetaCardis data custodians using the contact details provided in the repository. Analysis code will be made publicly available at https://github.com/kamoors/PPI_modelling_and_microbiomes upon publication of the peer-reviewed article and will be provided to editors and reviewers upon request during peer review.

## Acknowledgements

We acknowledge funding by the German Research Foundation to C.K. within the scope of CRC1182 (project A1.5), the Research Group miTarget (FOR5042) and the Cluster of Excellence ‘Precision Medicine in Chronic Inflammation’ (EXC2167) to C.K., A.F., and P.R. K.A.M. and this work were supported by the Add-on Fellowship of the Joachim Herz Foundation. M.A.G. gratefully acknowledges support from the Detlef Zillikens Clinician Scientist Academy of Precision Health in Schleswig-Holstein (PHSH) and the Clinician Scientist Academy [Kiel]. J.H received funding through the DFG (Project-ID 499552394 – CRC 1597). Microbiota sequencing was supported by the DFG Programme Core Facilities (project 568759757) and received infrastructure support from the DFG Excellence Cluster 2167 “Precision Medicine in Chronic Inflammation” (PMI) and the DFG Research Unit 5042 “miTarget”. This work was supported by the DFG Research Infrastructure NGS_CC (project number: 407495230) as part of the Next Generation Sequencing Competence Network (project number: 423957469). Short-read sequencing was conducted at the Competence Centre for Genomic Analysis (Kiel, Germany) supported by the state of Schleswig-Holstein. The project received funding from the European Union under the Horizon Europe grant agreement No. 101095470 (project miGut-Health).

## Supplementary materials

**Supplementary Figure S1.**
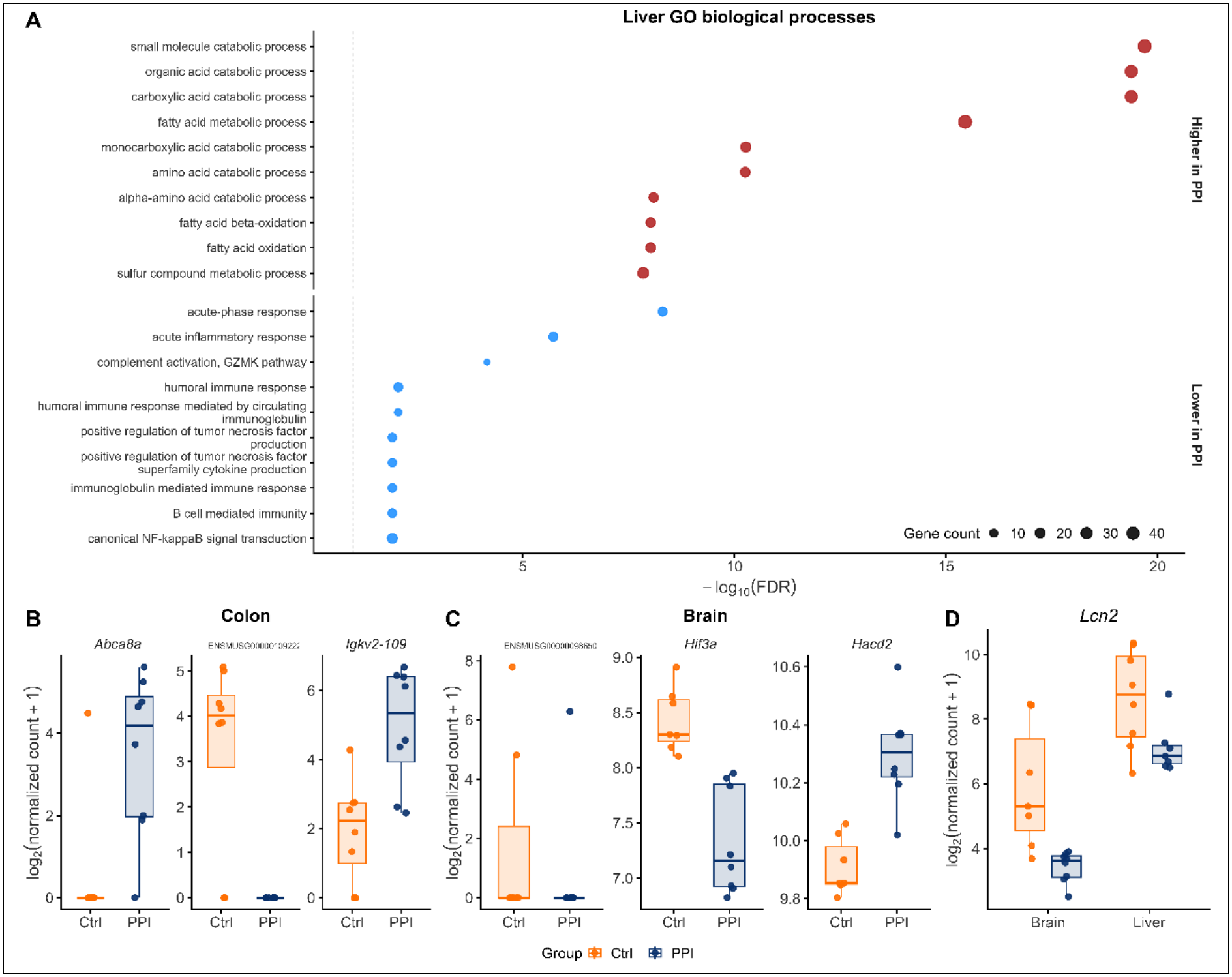
Liver Gene Ontology enrichment and expression of selected differentially expressed genes across tissues. **(A)** Gene Ontology biological-process terms enriched among liver genes with significantly higher or lower expression in recipients of PPI-associated microbiota (DESeq2 FDR < 0.1). The ten terms with the lowest enrichment FDR in each direction are shown. Point position indicates −log10(FDR), point size indicates gene count and colour denotes direction. The dashed line indicates FDR = 0.1. **(B,C)** Expression of genes differentially expressed in colon **(B)** and brain **(C)**. Boxplots show log₂(normalized counts + 1), with points representing individual mice. **(D)** Expression of *Lcn2*, which was differentially expressed in both brain and liver. Differential expression was tested separately within each tissue using DESeq2 on untransformed counts; all genes shown passed FDR < 0.1.

## Methods

### Cohort descriptions

The Kiel cohort consisted of 1,280 participants recruited through the PopGen (27) and FoCus (28) studies. Clinical metadata included demographic information, anthropometric measurements, medication use and microbiome profiling data. Within the cohort, 102 participants (8.0%) were classified as proton-pump inhibitor (PPI) users. The cohort included 795 female participants, of whom 62 (7.8%) were PPI users, and 485 male participants, of whom 40 (8.3%) were PPI users. The mean age of the cohort was 53.0 ± 14.6 years (median 54 years, interquartile range [IQR] 43– 64 years), while mean BMI was 30.7 ± 10.2 kg/m² (median 30.1 kg/m², IQR 22.6–36.3 kg/m²). PPI users were older and had higher BMI values compared with non-users. Medication burden was also higher among PPI users. Use of the Kiel cohort data was approved by the Kiel University Ethics Committee (A156/03 v. 29.7.2011 and A156/03 v. 29.7.2010).

Publicly available data from the MetaCardis cohort were used as an independent European validation cohort. MetaCardis is a multi-center study investigating relationships between the gut microbiome, host metabolism and cardiometabolic disease across several clinical populations (29). After quality-control filtering, the analysed dataset included 997 participants, including 147 PPI users (14.7%) and 850 non-users. The cohort contained 390 female participants, of whom 54 (139%) were PPI users, and 607 male participants, of whom 93 (15.3%) were PPI users. The mean age of the cohort was 57.7 ± 12.0 years (median 60 years, IQR 52–66 years), while mean BMI was 28.9 ± 7.7 kg/m² (median 27.0 kg/m², IQR 23.6–32.1 kg/m²). PPI users showed higher frequencies of diabetes, statin use, aspirin use and greater overall medication burden compared with non-users. Samples annotated as non-PPI users but showing detectable metabolomic signals for omeprazole, pantoprazole or dexlansoprazole were excluded prior to downstream analyses to reduce exposure misclassification.

### Cohort statistical analysis

Clinical metadata for the Kiel and MetaCardis cohorts included age, sex, body mass index (BMI), diabetes status, medication use and additional cohort-specific clinical variables. Medication burden was defined as the total number of broad medication classes used by each participant and was included as a covariate in downstream analyses. In the MetaCardis cohort, samples annotated as non-PPI users but showing detectable metabolomic signals for omeprazole, pantoprazole or dexlansoprazole were excluded to reduce exposure misclassification. Following this filtering step, metadata-defined PPI-use status was used as the exposure variable for all downstream analyses. Associations between PPI exposure and microbial, metabolic and model-derived molecular features were assessed using multivariable linear regression models adjusted for relevant demographic and clinical covariates. Shared covariates across cohorts included age, sex, BMI, diabetes status, medication burden, metformin use, statin use and aspirin use. Additional cohort-specific covariates, including nationality and creatinine in MetaCardis, were included where available. Associations were tested using multivariable linear models with heteroskedasticity-consistent standard errors calculated using the HC3 covariance estimator implemented in the sandwich R package (v3.1-1) (68, 69). Two-sided *P* values were adjusted using the Benjamini–Hochberg (BH) method, with FDR < 0.10 considered significant. Analyses were restricted to complete cases for all model covariates. To assess reproducibility across cohorts, cohort-specific effect estimates and standard errors from the Kiel and MetaCardis analyses were combined using fixed-effect inverse-variance weighted meta-analysis. Combined effect sizes, standard errors, z-statistics, p-values and BH-adjusted p-values were calculated for shared microbial, metabolic and reaction-level features. Cross-cohort agreement was further evaluated using Spearman correlation of cohort-specific effect estimates, concordance of effect directions and overlap of individually significant features.

### Bacterial abundance

For bacterial abundance analyses, compositional microbiome data were analysed using cohort-specific sequencing and statistical workflows. In the Kiel cohort, microbiome data in the form of 16S amplicon sequence data were generated in previous studies (27, 28). Briefly, following stool sample collection and DNA extraction, the V1–V2 region of the 16S rRNA gene was amplified using the primer pair 27F/338R and sequenced on the Illumina MiSeq platform after normalization and equimolar pooling of PCR products. Forward and reverse reads were merged using FLASH, and quality filtering was performed using UCHIME with exclusion of low-quality sequences based on Q-score thresholds. In the MetaCardis cohort, stool samples were collected and processed according to International Human Microbiome Standards (IHMS) guidelines as described previously (29). Total faecal DNA was extracted using standardized protocols and subjected to shotgun metagenomic sequencing using Ion Proton technology (Thermo Fisher Scientific). Sequencing reads underwent adapter trimming, quality filtering and removal of human and contaminant DNA prior to mapping against the integrated human gut microbial gene catalog. Taxonomic profiles were generated from metagenomic species-level abundance estimates following rarefaction and normalization procedures described previously (29). For differential abundance analyses, ALDEx2 (70-72) was applied in the Kiel cohort using centred log-ratio transformation, 256 Monte Carlo instances and generalized linear modelling, whereas MaAsLin2 (v1.18.0) (73) was used in MetaCardis with equivalent covariate adjustment and standard parameters. Features were filtered to retain taxa with non-zero variance and non-zero abundance in at least 10% of samples.

### Metabolic modelling

Microbiome community metabolic models were generated from bacterial abundance profiles using the AGORA2 genome-scale metabolic reconstruction database (22). Taxonomic assignments from 16S rRNA sequencing (Kiel cohort) and Metagenomic (MetaCardis) data were mapped to AGORA2 taxa using MARS (74). Some names in the dataset had to be adjusted due to changes in taxonomic identity. For the MetaCardis cohort, the AGORA2 model collection was extended to increase taxonomic coverage. Strain-level genome-scale metabolic models were reconstructed for 175 species not represented in AGORA2 through the DEMETER (75) where a whole genome sequence was available on NCBI Assembly. Curation against experimental data for named species was performed where possible. The reconstructions were validated through the DEMETER test suite to ensure biomass production, thermodynamic consistency, and agreement with available experimental data. These newly reconstructed models were combined with the existing AGORA2 strain-level models to generate pan-species models using the COBRA Toolbox. All reads for the mapped species were normalized per sample, so the resulting sum of relative abundances equalled 1. Species with relative abundances below 1e-6 were excluded and the dataset was renormalized to obtain the final relative abundance profiles. The MgPipe pipeline within the Microbiome Modelling Toolbox (23) implemented in the COBRA Toolbox (21) was used to construct microbiome community models from the relative abundance profiles and corresponding AGORA2 pan-species models. The EUAverageDietNew, defined in the COBRA Toolbox, was used to constrain the metabolic models and the microbiome biomass growth was confined to [1,1] mmol/day/person. Finally, the microbiome community models were integrated with the corresponding male or female Whole-Body Human Models (WBMs) Harvey and Harvetta (26) using the combineHarveyMicrobiota function in the COBRA Toolbox.

AGORA2 strain-level oxygen-requirement annotations were used to classify reactions according to the oxygen-tolerance profiles of the species carrying them. Aerobic strains were classified as aerotolerant, facultative anaerobes and aerotolerant strains as facultative, and obligate anaerobes as anaerobic. For species represented by at least two strains with recognized annotations, category scores were calculated as the proportions of annotated strains assigned to each category. Reaction frequency within each species was defined as the proportion of its strain models containing the reaction, with absent reactions assigned a frequency of zero. For reactions present in at least three species and showing between-species variation, Spearman correlations were calculated across 347 species between reaction frequency and each oxygen-tolerance score. A reaction was assigned to the category with the largest correlation when this correlation was at least 0.20 and exceeded the second-largest correlation by at least 0.20; all other reactions were classified as mixed. These classes describe the ecological distribution of reactions across species models rather than their biochemical oxygen requirements. Reaction-level meta-analysis effect estimates were compared among classes using all six pairwise two-sided Wilcoxon rank-sum tests, with P values adjusted across the six comparisons using the Benjamini–Hochberg procedure. Robustness was assessed across all combinations of correlation and separation thresholds ranging from 0.10 to 0.30 in increments of 0.05.

Microbiome community models were used to calculate MgPipe net maximal production capabilities, hereafter termed net secretion, for all metabolites that could be secreted by at least one community member. Under *EUAverageDietNew* dietary constraints and with community biomass growth fixed at [1,1] mmol/day/person to enable comparison across microbiomes, flux variability analysis separately determined the maximal feasible faecal secretion and dietary uptake of each metabolite. Net secretion was calculated as the absolute value of the maximal faecal secretion flux plus the minimal dietary exchange flux, with uptake represented by a negative flux. For model-derived net secretion fluxes, missing values were set to zero and features with zero variance were removed prior to analysis. Since flux distributions were right-skewed, response variables were transformed using log1p before modelling. Reaction abundance profiles representing the abundance of each metabolic reaction within each microbial community were analysed using the same regression framework. Reaction abundances were calculated using the MgPipe fastCalculateReactionAbundance function. For each reaction, its presence or absence was determined from the reaction content of each species-level pan-model, and its abundance in a sample was calculated as the sum of the relative abundances of all species whose models contained that reaction. Reaction abundance values therefore ranged from zero to one and represent the abundance-weighted genomic potential for a reaction rather than its predicted flux or activity. Missing values were set to zero, reactions with zero variance were removed, and values were log-transformed after adding a pseudocount of 10^-6^. The sum of all reaction-abundance values within each sample was included as a covariate to account for differences in the overall abundance-weighted reaction repertoire. Exploratory subsystem over-representation analysis was performed separately for reactions with positive and negative meta-analysis effects passing BH-adjusted *P* < 0.10. One-sided hypergeometric tests implemented in clusterProfiler (76) were conducted using all tested reactions with AGORA2 subsystem annotations as the background, and enrichment *P* values were adjusted using the Benjamini–Hochberg procedure. Subsystems containing fewer than five tested reactions were excluded. Because reaction abundances are correlated through shared species and metabolic networks, these results were interpreted as descriptive summaries of the reaction-level associations rather than independent confirmatory evidence.

To assess blood-accessible metabolic potential in the combined community microbiome– whole-body models, maximum feasible fluxes were calculated for a predefined panel of 323 blood-compartment metabolites. For each metabolite an artificial demand reaction consuming the metabolite from the blood compartment (DM_*[bc]) was added and maximized independently. The resulting fluxes, expressed in mmol/day/person, represent the maximum model-feasible supply of each metabolite to the blood compartment rather than predicted circulating concentrations. Fluxes were rounded to 10^-16^ and features with a fraction of unique flux values below 0.1 were excluded. Associations with PPI exposure were tested using the cohort-specific regression models and HC3 robust standard errors described above, followed by the same cross-cohort meta-analysis and false-discovery-rate control. All simulations used the combined microbiome–host models.

### Human samples

Faecal donors were selected from the Institute of Transfusion Medicine (ITM) blood-donor cohort at UKSH, Kiel, comprising participants from Northern Germany who provided stool samples and dietary, lifestyle and medical-history questionnaires, as described previously (77). Samples were initially selected based on proton pump inhibitor (PPI) use. The initial cohort comprised four PPI-treated individuals, including three pantoprazole users and one omeprazole user. For each PPI-treated individual, a matched control was selected based on sex and year of birth, resulting in an initial matched set of PPI users and controls. To ensure consistency of PPI-type taken within the treatment group, the individual treated with omeprazole and the corresponding matched control were excluded from further analyses. The final study population therefore consisted of six individuals, including three pantoprazole-treated patients and three matched controls. Donor characteristics and recorded diagnoses are provided in Table 1. This matched case–control design was used to minimize confounding by age and sex while maintaining a homogeneous exposure group with respect to PPI treatment. The study was conducted in accordance with the Declaration of Helsinki, and ethical approval was granted by the ethics committee at Kiel University (D590/22, relating to A103/14).

### Ex vivo Anaerobic Incubations of Human Stool

Stool samples were obtained from six human donors, including three proton pump inhibitor (PPI) users and three non-users (control group). Samples were stored in glycerol at −80 °C until use. *Ex vivo* cultivation was performed in prereduced mGAM medium (HiMedia, M2079; prepared according to the manufacturer’s instructions) for 96 hours, sampled at five time points under strictly anaerobic conditions in an anaerobic chamber (Coy Laboratory Products) maintained with a gas mixture of 5 % H₂, 10 % CO₂, and 85 % N₂. Media were prereduced inside the chamber prior to inoculation. Cultures were incubated at 37 °C with gentle shaking (200 rpm) using an integrated incubator. On Day 0, 10 µL of glycerol-preserved stool was inoculated into 15 mL cell culture tubes (Greiner Bio-One, 188241) containing 10 mL prereduced mGAM medium. Samples were collected at five defined time points during the 96-hour experiment (0, 24, 48, 72, and 96 h post-incubation) to monitor temporal changes in microbial composition and activity. At each time point, cultures were vortexed thoroughly, and 2 mL of culture was removed and replaced with 2 mL of fresh prereduced mGAM medium. From the collected sample, 1 mL was centrifuged at 14.000 rpm for 10 min at 4 °C (Eppendorf 5425R centrifuge). The supernatant was stored at −80 °C for metabolomic analysis, and the corresponding cell pellet was preserved at −80 °C for 16S rRNA gene sequencing. Sterile mGAM medium was included as a negative control. Optical density at 600 nm (OD₆₀₀) was measured daily from an aliquot of the 2 mL sample collected at each time point using a Tecan Infinite M Nano+ spectrophotometer to monitor bacterial growth.

### Gnotobiotic Animal Experimentation

All animal experiments were performed according to approved animal protocols approved by the local animal safety review board the Ministry of Agriculture, Rural Areas, European Affairs and Consumer Protection (MLLEV) of Schleswig Holstein and conducted in accordance with FELASA guidelines and German animal welfare law (IX554-62160/2024(48-6/24)). Germ-free (GF) C57BL/6J male mice aged approximately 15 weeks (n=16) were initially imported from the gnotobiotic mouse unit of the Hannover Medical School and then bred inhouse at the Central Animal Facility (ZTH) of the University Hospital Schleswig Holstein (UKSH, Kiel, Germany) under aseptic conditions in sterile isolators (Class biologically clean, Wisconsin, USA). All mice were held under a 12h light/12h dark cycle at 22°C (±2°C) and a humidity of 55-60%. Mice had access to autoclaved mouse breeding food (1318, hybrid pellet of diet 1310, Altromin, Lage, Germany) and sterile water *ad libitum*. Mice were aseptically extracted from isolators and handled under a sterile flow hood for weighing and stool sample collection at the beginning of the experiment. Two to three germ-free mice per donor were then colonized by faecal microbiota transfer as described previously (66, 78-80). Briefly, recipient germ-free mice were orally gavaged with 200µl of cryoprotected stored human faecal material resuspended in sterile PBS with 20% glycerol (∼75 mg faecal material per ml) using a 18G reusable feeding needle (Fine Science Tools, Heidelberg, Germany) immediately upon thawing. Colonized mice were then transferred into sterile gnotocages (IsoCage, Tecniplast, Italy) and housed under gnotobiotic routines for the experimental duration. Mice were extracted from the gnotocages under aseptic conditions once a week for weighing and faecal collection. The sterile flow hood and all handling materials were disinfected between each mouse group. After two weeks, all mice were sacrificed by cervical dislocation for tissue and fluid collection.

### 1cS rRNA gene sequencing of ex vivo pellets and mouse faecal samples

Paired end 16S rRNA gene sequences were processed in R with the dada2 R package (version 1.32.0). Read quality profiles were inspected to guide filtering and truncation. Forward reads were truncated at 240 bases and reverse reads at 190 bases, reads with ambiguous bases were discarded, maximum expected errors were limited to two for each read direction, bases with quality below two triggered truncation, and phiX reads were removed. Error models were learned separately for forward and reverse reads and dereplication was performed prior to denoising. Denoised reads were merged by sample to reconstruct amplicon sequence variants, and a consensus method was used to remove chimeras. Read counts were tracked across all steps to document retention, including the ratio of non chimeric to merged reads. A mock community sample was used as a sanity check of the inferred sequence set. Taxonomic annotation was performed against the SILVA v138.2 training set using DADA2, and species level labels were added when available.

For downstream community analyses, ASV counts were converted to relative abundances within each sample. Bray–Curtis dissimilarities were calculated using phyloseq (v1.54.2) (81). To assess preservation of donor-specific community composition following ex vivo cultivation and gnotobiotic transfer, the Bray–Curtis dissimilarity between each original donor sample and samples derived from the same donor was compared with its dissimilarity to samples derived from the other donor microbiotas. For the ex vivo analysis, day 0 samples were excluded and measurements from culture days 1–4 were retained. Recipient and non-recipient Bray–Curtis dissimilarities were compared separately for each donor using Wilcoxon rank-sum tests. Community composition was additionally visualized using non-metric multidimensional scaling (NMDS) based on Bray–Curtis dissimilarities calculated from relative-abundance-transformed ASV profiles. ASV counts, taxonomic annotations, genus- and species-level abundance tables, sample metadata and Bray–Curtis source data are provided in Supplementary Data 2.

### RNA-seq: Alignment, quality control, and differential expression

Raw paired-end RNA-seq reads were quality controlled using MultiQC (v1.30) (82) and trimmed using fastp (v0.23.4) (83). Pre-processed reads were quantified at the transcript level using Salmon (v1.10.2) in quasi-mapping mode. A Salmon index was built from the Mus musculus GRCm39 reference genome (Ensembl release 112) by concatenating transcript sequences (cDNA) with the genome to generate a gentrome and providing the corresponding genomic decoy list. Transcript-level abundance estimates were summarised to the gene level using tximport (v1.34.0) (84) together with the corresponding Ensembl transcript-to-gene annotation. Transcriptomic outliers were identified independently within each organ using variance-stabilized expression data generated with the variance stabilizing transformation implemented in DESeq2 (85). Multiple complementary metrics were evaluated, including sample-to-sample correlation structure, correlation with the corresponding group centroid, median Cook’s distance across genes, and multivariate Mahalanobis distance calculated from the first five principal components. Samples were ranked across metrics and combined into an overall outlier score. One liver sample (25Jul1866-C02) and one brain sample (25Jul1866-G04) consistently showed the strongest evidence of atypical transcriptomic profiles and were excluded from downstream analyses.

Genes were retained when they had at least ten counts in a minimum of 20% of samples within the corresponding organ-specific dataset. Differential expression was assessed independently within each organ using DESeq2 (v1.44.0) (85). Separate DESeqDataSet objects were generated using DESeqDataSetFromTximport, with experimental group specified as the design factor (∼ group) and control recipients used as the reference group. PPI-versus-control differences were assessed using Wald tests, and multiple testing was controlled using the Benjamini–Hochberg procedure. Genes with FDR < 0.10 were considered differentially expressed. Since multiple recipient mice received microbiota originating from the same human donor, complementary analyses were performed using the limma-voom framework (86, 87) to assess the influence of donor-level correlation. Following normalization and voom transformation, residual correlation among mice receiving microbiota from the same donor was estimated using duplicateCorrelation, and donor identity was included as the blocking factor during model fitting. Organ-specific PPI-versus-control contrasts were evaluated using moderated t-statistics with Benjamini–Hochberg correction. A matched limma-voom model without donor blocking or duplicateCorrelation was additionally fitted to assess the effect of explicitly modelling the donor-level correlation structure. Further sensitivity analyses evaluated the potential influence of measured mouse-level and housing-related covariates. In DESeq2, additional models included recipient age (∼ age_z + group), isolator (∼ isolator + group), or both (∼ age_z + isolator + group). Corresponding donor-blocked limma-voom models additionally considered age, isolator, age together with isolator, and age, isolator and cage position. These variables were treated as sensitivity covariates rather than primary adjustment factors because recipient mice receiving microbiota from the same human donor were intentionally housed together and age matched as far as experimentally feasible, resulting in substantial alignment between donor identity, housing variables and recipient age. The alternative models were therefore used to assess robustness of the group contrast rather than to define the primary differential-expression results.

Gene-set enrichment analysis was performed separately for each organ using a preranked strategy implemented with fgsea (v1.36.2) (88). Ensembl identifiers were converted to gene symbols using AnnotationDbi (v1.66.0) and org.Mm.eg.db (v3.19.1). Genes were ranked according to the DESeq2 Wald statistic. Where multiple Ensembl identifiers mapped to the same gene symbol, the entry with the largest absolute Wald statistic was retained. Hallmark and Reactome pathway definitions were obtained from the Molecular Signatures Database using msigdbr (v26.1.0) (89). Gene sets were restricted to pathways containing 15–300 genes for Hallmark analyses and 15–500 genes for Reactome analyses. Enrichment was assessed using fgseaMultilevel, yielding enrichment scores, normalized enrichment scores (NES), nominal P values, FDR-adjusted P values and leading-edge genes. Multiple testing across pathways was controlled using the Benjamini–Hochberg procedure. Directional Gene Ontology Biological Process over-representation analysis was additionally performed for liver, which was the only organ yielding sufficient significant genes for meaningful enrichment analysis. Significant liver genes (FDR < 0.10) were separated according to the direction of the PPI-versus-control effect and analysed independently using enrichGO in clusterProfiler (76). Ensembl identifiers were mapped to Entrez identifiers using org.Mm.eg.db, and all genes tested in the corresponding differential-expression analysis were used as the background universe. Benjamini–Hochberg correction was applied across tested GO terms, with FDR < 0.10 considered significant. The donor-aware limma-voom gene set was analysed in parallel as a robustness comparison.

### Nuclear Magnetic Resonance (NMR)-Based Non-Targeted Metabolomics

Bacterial supernatant sample preparation consisted of mixing 180 µL of supernatant with 20 µL of NMR buffer (100% D2O, 1.5 M phosphate buffer with 0.1% trimethylsilyl-tetradeuteropropionic acid (TSP), pH 7.4). To extract the aqueous metabolites of caecal content, we homogenized 50 mg sample in 1 mL H2O using ceramic beads (NucleoSpin, Macherey–Nagel, Dueren, Germany) and a TissueLyser (Qiagen, Hilden, Germany) mixing the sample for 3 x 20 sec at 5000 rpm with a 10 sec cooling break (< 0°C). Subsequently, the homogenate was centrifuged (13.000 rpm for 10 mins at 4°C), the supernatant evaporated with a SpeedVac, the dried extract reconstituted in 180 µL H2O and mixed with 20 µL NMR buffer. Samples were transferred to 3 mm NMR tubes. NMR analysis happened immediately (autosampler at 4°C) and in a randomized order. NMR spectra were measured with a Bruker 800 MHz spectrometer operating at 800.35 MHz. A standard one-dimensional pulse sequence (noesygppr1d) delivered an overview of all molecules. The acquisition parameters were as follows: water suppression irradiation during recycle delay (2 s), mixing time = 200 ms, 90◦ pulse = 12.5 µs. We collected 512 scans into 64 K data points with a spectral width of 12 ppm. The software TopSpin 3.7 (Bruker BioSpin, Ettlingen, Germany) was used for processing, i.e., Fourier transformation, manual phasing, baseline correction and calibration to TSP (δ 0.00). Data were imported into Matlab software R2024b (Mathworks, Natick, MA, USA) and further processed, i.e., the water region removed, baseline adjusted (90), peaks aligned (91) and spectra normalized (92). Identification and relative quantification of metabolites was done using the workflow described in Heinzmann *et al.* using a peak picking algorithm and subsequent clustering of peaks based on correlation and hierarchical clustering analysis approaches and supported by subsequent series of 2-dimensional NMR analyses (93).

### Metabolomics of bacterial supernatants using HILIC UHPLC-MS/MS

An aliquot of 250 µL of each bacterial supernatant was mixed on a laboratory vortex mixer for 10 seconds and transferred to 750 µL acetonitrile (ACN; LiChrosolv, hypergrade for LC–MS; Merck KGaA), which was spiked with L-Leucine 5,5,5-d3 at 5 mg/L (99 atom % D; Sigma-Aldrich). The mixture was mixed again for 10 seconds on a laboratory vortex mixer and centrifuged for 10 minutes at 4 °C and 13100 rpm. Samples were transferred into sterile tubes and evaporated at 40 °C with a SpeedVac concentrator (Savant SPD121P; Thermo Fisher Scientific, Waltham, MA, USA) and reconstituted in 1 mL of pure milli-Q water (Milli-Q Integral Water Purification System; Billerica, MA, USA). The samples were analyzed by using a UHPLC system (Acquity; Waters, Eschborn, Germany) coupled to a quadrupole time-of-flight (TOF) mass spectrometer (maXis; Bruker Daltonics, Bremen, Germany), as described previously (94).

Ammonium acetate (NH4Ac; LiChropur eluent additive for LC–MS; Merck KGaA) at 0.5 mol/L was adjusted to pH 4.6 with glacial acetic acid (Honeywell; Fluka, Seelze, Germany). Milli-Q water was obtained from a Milli-Q Integral Water Purification System (Billerica, MA, USA). Polar metabolites were separated by HILIC by using an iHILIC-Fusion UHPLC column SS (100 × 2.1 mm, 1.8 μm, 100 Å; HILICON AB, Umea, Sweden). The eluent compositions were as follows: Eluent A consisted of 5 mmol/L NH4Ac (pH 4.6) in 95% ACN (pH 4.6), and eluent B consisted of 25 mmol/L NH4Ac (pH 4.6) in 30% ACN. We started with 0.1% B, keeping it constant for two minutes, then increased B to 99.9% over 7.5 minutes. The condition of 99.9% B was kept for two minutes and reversed to 0.1% B within 0.1 minutes, held for 0.1 minutes. The run was completed after 12.1 minutes, and the column was equilibrated for five minutes before the next injection. The flow rate was set to 0.5 mL/min, the column temperature to 40 °C, and the sample manager was cooled to 4 °C; 5 μL of the sample was injected into the column (partial loop). The weak and strong washes consisted of 95% and 10% ACN, respectively.

Mass spectra were acquired at electrospray ionization (ESI) of positive and negative modes (+/−). Spectrometric data were acquired in line and profile mode with an acquisition rate of 5 Hz from 50 to 1500 Da. Fragmentation experiments were set to the data-dependent mode (MS/MS [Auto]), where the three most intense ions were fragmented within one scan when a count reached over 2000. Ions were excluded after acquiring three MS/MS and reconsidered for fragmentation after six seconds. The collision energy was set to 20 eV for (+/−) mode with an isolation width of 8 Dalton. The electrospray ionization source parameters were as follows: capillary voltage of 4500 V for (+) and 4000 V for (−), end plate offset of (+/−) 500 V, nebulizer gas of 2 bar, dry gas of 10 L/min, and dry heater of 200 °C. Before measurements, the MS was calibrated using the ESI-L Low Concentration Tuning Mix (Agilent, Santa Clara, CA, USA). The ESI-L Low Concentration Tuning Mix (diluted 1:4 [v/v] with 75% ACN) was injected in the first 0.3 minutes of each run by a switching valve for internal recalibration by post-processing software.

### Metabolomics of serum, caecal and faecal content using RP UHPLC-MS/MS

Serum samples were thawed (room temperature) and 50 µL were transferred into 150 µL ice-cooled methanol, containing 0.1 mg/L d4-glycodeoxycholic acid (Glycodeoxycholic-2,2,4,4-d4 acid, ≥98 atom % D, ≥98% (CP), Sigma-Aldrich). Samples were shaked for 30 seconds on a vortexer and centrifuged for 10 minutes at 14000 rpm and 4°C. Supernatant was transferred into sterile tubes until analysis. Caecal content (x̄ = 28 mg) and faecal pellets (x̄ = 22 mg) were transferred in sterile ceramic bead tubes (NucleoSpin Bead Tubes; Macherey-Nagel, Dueren, Germany) and extracted with 1 mL of chilled methanol (−20 °C; LiChrosolv, Supelco; Merck KGaA, Darmstadt, Germany) containing 0.1 mg/L d5-taurocholic acid (Taurocholic Acid-d5 Sodium Salt, Toronto research chemicals). Caecal and faecal matter was homogenized and extracted with a Precellys Evolution Homogenizer (Bertin Corp., Rockville, MD, USA; 4,500 rpm, three 40-second cycles with a two-second pause). The samples were centrifuged for 10 minutes at 12000 rpm and 4°C, and the supernatant was transferred into sterile tubes until analysis.

Samples were measured on the same UHPLC-MS/MS system as above but opted for reversed-phase chromatography (RP) using an octyl reversed-phase column (C8: 1.7 μm, 2.1 × 150 mm, Acquity™ UPLC BEH™) (95). The MS parameters were as described above. Only the collision energy tandem MS experiments differed and was set to 20 eV in (-) and 30 eV in (+) mode.

The eluent compositions were as follows: Eluent A was 5 mM NH4Ac and 0.1% acetic acid in water and eluent B was pure ACN. A gradient profile was applied by starting at 10% B for 1 minute, increasing to 22% B within 2 minutes, followed by an increase to 27% B within 4 minutes, taken over by a linear gradient to 95% B within 13 minutes. The 95% B was held for 2.5 minutes, returning to initial 10% B within 2.5 minutes, resulting in a total run time of 25 minutes. Prior to next measurement, a pre-run time of 2 minutes was applied to re-equilibrate the column. The flow rate was set to 0.35 mL/min, the column temperature to 60°C, and the sample manager was cooled to 4°C; 5 μL of the sample was injected into the column (partial loop). A mixture of isopropanol:acetonitrile:methanol:water (1:1:1:1, v/v) as strong needle wash and an acetonitrile:water (9:1, v/v) mixture as weak needle and seal wash was used.

### Metabolomic data processing and metabolite identification

Raw data was calibrated by the automatic internal calibration (Tuning Mix ES-TOF (ESI) List for -/+ mode) in quadratic mode with search range of 0.2 *m*/*z* and intensity threshold of 500 and exported automatically to .mzML files. Generation of feature tables and metabolite identification was done in Mzmine version 4.5.0 (96). Experimental and in silico spectral libraries were downloaded from different sources including MassBank of North America, Global Natural Products Social (97) and MS-DIAL LipidBlast (version 69) (98). We also used Sirius to facilitate annotation of MS/MS, and we considered only features with ConfidenceScore > 0.7 (99). We ranked features following, metabolites matched by authentic standards were given Rank 1, metabolites identified by spectral match as Rank 2 and by Sirius as Rank 3. Non-resolved isomers are indicated by slash (/) in figures. Furthermore, some metabolites were identified visually and ranked as 1: 3-Hydroxy Stearic Acid was identified based on this work (100). Other 3-hydroxy fatty acids (3-Hydroxydecanoic acid, 3-Hydroxydodecanoic acid, 3-Hydroxytetradecenoic acid, FA 14:2;O-3OH, 3-Hydroxytetradecanoic acid) were identified based on a loss of acetic acid fragment in (-) MS/MS (101). Mirror plots of lysophospholipids (LPC, LPE, LPI) and acylcarnitines (CAR) were manually checked in Mzmine. 10-Hydroxy Stearic Acid was identified based on this work (102). Stercobilin was identified based on this work (51). Mesobilirubinogen was identified based on the Spectrum of https://mona.fiehnlab.ucdavis.edu/spectra/display/MT000068. Data was normalized to peak areas of internal standards (Bacterial supernatants: L-Leucine 5,5,5-d3, serum: d4-glycodeoxycholic acid, and caecum/faeces: d5-taurocholic acid) and weight of caecal content and faecal pellets.

### Statistical Analysis

LC-MS and NMR caecum/faeces datasets, serum LC-MS, as well as HILIC ex vivo datasets, were analysed using a unified statistical framework. Blank and quality-control samples were excluded where applicable, and analyses were restricted to annotated metabolites. LC–MS annotations were manually curated, and duplicated features were resolved without reference to treatment effects. Features detected in both ionization modes were considered to represent the same analyte when their corrected metabolite names matched and their retention times differed by no more than 0.10 min. For metabolite-level analyses, one representative feature was retained per curated metabolite based on detection frequency, annotation confidence and analytical performance; unresolved retention-time-separated features were retained as distinct analytes where appropriate. LC–MS intensities were transformed as log_2_(*x* + *p*), where *p*was half the smallest positive intensity observed for the corresponding feature. Metabolites with insufficient detection or no measurable variance were excluded. For NMR data, occasional negative baseline-corrected peak areas were set to zero before applying the same feature-specific log transformation. Resonances assigned to the same metabolite were standardized across samples and combined using their median standardized value, allowing differently scaled resonances to contribute comparably. Ex vivo analyses included culture days 1–4; day 0 measurements were used for trajectory analyses but excluded from differential testing.

Differential abundance was assessed using linear models implemented in limma. All recipient mice or culture measurements were retained, and the correlation among samples derived from the same microbiome donor was estimated using duplicateCorrelation and incorporated into model fitting. Mouse serum and caecal NMR datasets were modelled as a function of microbiome treatment group. Caecal and faecal LC–MS data were analysed jointly using a model containing treatment group and compartment, with additional compartment-specific and group-by-compartment analyses. Ex vivo NMR and HILIC data were modelled using treatment group and categorical culture day, with donor included as the blocking factor. Models were fitted using robust empirical-Bayes moderation. Positive coefficients indicate higher abundance in samples receiving or cultured with PPI-associated microbiomes. Effect estimates are reported as log2 fold changes for LC–MS features and as differences in standardized metabolite scores for grouped NMR resonances, together with 95% confidence intervals. P values were adjusted using the Benjamini–Hochberg procedure within each dataset and contrast, and FDR < 0.10 was considered significant. PCA was used for exploratory visualization of global metabolomic variation using log-transformed and scaled intensities. Ordinations were interpreted descriptively given the clustering of samples within six donor microbiomes.

### Cross-layer association analyses

Genus–metabolite associations were evaluated separately within each metabolomics dataset using rank-standardized linear models adjusted for microbiome treatment group. Models for the ex vivo cultures additionally included categorical culture day and accounted for repeated measurements originating from the same inoculum donor. Effect estimates therefore represent the change in rank-standardized genus CLR abundance associated with metabolite abundance after adjustment for the experimental design. P values were corrected using the Benjamini– Hochberg procedure within each microbiome–metabolomics analysis family, and associations with FDR < 0.1 were considered significant. Robustness was assessed by repeating each significant association after sequential exclusion of each microbiome donor.

Metabolite–gene associations were analysed separately for each metabolite and tissue using DESeq2. Metabolite concentrations were standardized to a mean of zero and unit variance, and RNA-seq counts were modelled using the design ∼ Group + Metabolite. The metabolite coefficient therefore represents the gene-expression association per one-standard-deviation increase in metabolite abundance after adjustment for the mean difference between recipients of control- and PPI-associated microbiomes. The analysis included differentially altered serum, caecal, faecal RP LC–MS and caecal NMR metabolites selected at metabolomics FDR < 0.1. For caecal and faecal RP LC–MS, metabolites were selected from the combined compartment-adjusted analysis and subsequently modelled using their compartment-specific abundance measurements.

Gene-level P values were adjusted by the Benjamini–Hochberg method both within each metabolite-specific model and across all gene–metabolite tests within each metabolomics-layer–tissue family. Pathway enrichment was assessed using preranked gene set enrichment analysis based on the signed DESeq2 Wald statistic for the metabolite coefficient. Directional over-representation analysis was additionally performed separately for positively and negatively associated genes passing within-metabolite FDR < 0.1, using all genes tested in the corresponding model as the background. Mouse-native Hallmark and Reactome gene sets were used, and pathway P values were corrected within the respective metabolite, gene-set library and analysis family.

